# Dual RNA-seq study of the dynamics of coding and non-coding RNAs expression during *Clostridioides difficile* infection in a mouse model

**DOI:** 10.1101/2024.06.28.601227

**Authors:** Victor Kreis, Claire Toffano-Nioche, Cécile Denève-Larrazet, Jean-Christophe Marvaud, Julian R Garneau, Florent Dumont, Erwin L van Dijk, Yan Jaszczyszyn, Anaïs Boutserin, Francesca D’Angelo, Daniel Gautheret, Imad Kansau, Claire Janoir, Olga Soutourina

## Abstract

*Clostridioides difficile* is the leading cause of healthcare associated diarrhoea in industrialized countries. Many questions remain to be answered about the mechanisms governing its interaction with the host during infection. Non-coding RNAs (ncRNAs) contribute to shaping virulence in many pathogens and modulate host responses, however, their role in *C. difficile* infection (CDI) has not been explored. To better understand the dynamics of ncRNAs expression contributing to *C. difficile* infectious cycle and host response, we used a dual RNA-seq approach in a conventional murine model. From the pathogen side, this transcriptomic analysis revealed the upregulation of virulence factors, metabolism and sporulation genes, as well as the identification of 61 ncRNAs differentially expressed during infection that correlated with the analysis of available raw RNA-seq datasets from two independent studies. From these data we identified 118 potential new transcripts in *C. difficile* including 106 new ncRNA genes. From the host side, we observed the induction of several pro-inflammatory pathways and, among the 185 differentially expressed ncRNAs, the overexpression of microRNAs (miRNAs) previously associated to inflammatory responses or unknown long ncRNAs and miRNAs. A particular host gene expression profile could be associated to the symptomatic infection. In accordance, the metatranscriptomic analysis revealed specific microbiota changes accompanying CDI and specific species associated with symptomatic infection in mice. This first adaptation of *in vivo* dual RNA-seq to *C. difficile* contributes to unravelling the regulatory networks involved in *C. difficile* infectious cycle and host response and provides valuable resources for further studies of RNA-based mechanisms during CDI.

**Importance:** *Clostridioides difficile* is a major cause of nosocomial infections associated with antibiotic therapy classified as an urgent antibiotic resistance threat. This pathogen interacts with host and gut microbial communities during infection, but the mechanisms of these interactions remain largely to be uncovered. Noncoding RNAs contribute to bacterial virulence and host responses, but their expression has not been explored during *C. difficile* infection. We took advantage of the conventional mouse model of *C. difficile* infection to look simultaneously to the dynamics of gene expression in pathogen, its host and gut microbiota composition providing valuable resources for future studies. We identified a number of ncRNAs that could mediate the adaptation of *C. difficile* inside the host and the crosstalk with the host immune response. Promising inflammation markers and potential therapeutic targets emerged from this work open new directions for RNA-based and microbiota-modulatory strategies to improve the efficiency of *C. difficile* infection treatments.

## Introduction

*Clostridioides difficile* is an anaerobic spore-forming bacterium and the major cause of nosocomial infections associated with antibiotic therapy [1]. The major risk factors to contract *C. difficile* infections (CDI) are advanced age, the use of broad-spectrum antibiotics and immune system deficiencies. The disruption of the colonic microbiota by antimicrobial treatments precipitates colonization of the intestinal tract by *C. difficile* and ultimately leads to infection. Increasing severe forms and high recurrence rates favoured by persistent dysbiosis motivate the studies of *C. difficile* pathogenesis to develop synergistic and alternative treatments of CDI. Several *C. difficile* virulence factors have been identified, the toxins mainly responsible for epithelium lesions and clinical signs, as well as colonization factors like flagella and surface proteins. However, many aspects of *C. difficile* pathogenesis control remain poorly understood [2]. Several bacterial factors, in particular, *C. difficile* toxins and flagella, have been described to activate the inflammatory response [3, 4], which aims to clear the pathogen but can also contribute to the severity of intestinal lesions through an uncontrolled inflammatory process. Better understanding the regulations of both host response and the bacterial virulence factors expression during the infection is essential to improve our understanding of this important human pathogen.

During infection, bacteria reprogram the expression of their genes in response to diverse environmental constraints. Intensive studies of bacterial transcriptomes have shown the presence of a large number of non-coding RNAs (ncRNAs) [5] participating in the regulation of adaptive and pathogenic processes [6, 7]. Like in other pathogens, regulatory RNAs may shape virulence of *C. difficile.* Bioinformatics, RNA-seq and genome-wide promoter mapping identified more than 200 ncRNAs of different functional classes in *C. difficile* suggesting the diversity of RNA-based mechanisms for successful development of *C. difficile* inside the host [8–10].

Among them, several riboswitches responding to the signalling molecule c-di-GMP, co- ordinately control motility and biofilm formation, while multiple CRISPR (<u>c</u>lustered regularly interspaced <u>s</u>hort <u>p</u>alindromic repeats) RNAs are expressed to provide efficient defence against foreign genetic invaders for *C. difficile* survival in phage-rich gut communities [9, 11–13]. Antisense RNAs act as antitoxins within type I toxin-antitoxin modules contributing to prophage stability [14–16] and *trans*-acting ncRNAs work in concert with the RNA chaperone protein Hfq to control the metabolic adaptations, biofilm formation, stress responses and sporulation [17–19].

From the host side, ncRNAs including microRNAs (miRNAs) and long noncoding RNAs (lncRNAs) have been largely involved in the regulation of host inflammatory response and outcome of the infectious diseases [20]. In general, ncRNAs and, in particular, the miRNAs, operate in a complex network. A global view of differentially expressed ncRNAs in the host during CDI is currently missing. Since the modulation of the host response impacts dramatically the clinical outcome [4], deciphering the role of the host ncRNAs will provide new perspectives to control severe forms and recurrences of CDI.

We used here a dual RNA-seq [21] for simultaneous monitoring of the host responses to infection and bacterial riboregulators involved in host-pathogen interactions, successfully adapted to various infection models [22–25]. *In vivo* transcriptomics have been analysed separately from pathogen or host side in several studies using microarrays or RNA-seq during CDI [26–29]. The first *in vivo C. difficile* transcriptomic studies have been performed in a mono-associated mouse model of CDI [26, 29, 30] and the use of microarrays excluded the ncRNAs from this analysis. We explore here for the first time the dual transcriptome in a conventional *in vivo* model of CDI that better mimics the human infection to identify novel ncRNAs shaping *C. difficile* virulence and host response. As expected, several *C. difficile* virulence markers and host inflammatory response genes were induced during infection. Our dual RNA-seq analysis identified 61 ncRNAs among differentially expressed genes in *C. difficile in vivo* that could mediate the adaptation of *C. difficile* inside the host. From the host side, 185 ncRNAs were differentially expressed during infection including numerous lncRNAs and miRNAs enriching the regulatory network governing host response to pathogen infection. A particular gene expression profile from the host was associated with symptomatic CDI paving the way for a better understanding of the process leading from colonisation to symptomatic infection.

## Material and Methods

### Bacterial strains, growth conditions and preparation of spores

This work was performed with the *C. difficile* strain 630Δ*erm* [31], derived from the clinical 630 strain isolated from a patient suffering a pseudomembranous colitis, widely used for ncRNA studies and mouse model experiments [27, 30, 32]. For the *in vitro* culture, vegetative *C. difficile* cells was grown in Brain Heart Infusion (BHI) at 37°C in an anaerobic chamber (5% H2, 5% CO2, 90% N2, Jacomex, France) during 8h to reach late exponential growth phase (OD600 around 1.5). For the mouse challenge, *C. difficile* spores were prepared as previously described [33]. Vegetative cells were eliminated by heating at 70°C during 25 min and spores were numerated on BHI solid medium supplemented with taurocholate (0.1%) incubated 48h at 37°C under anaerobic conditions.

### Animal model and treatment

All animal assays were conducted in accordance with the institutional guidelines that follow the European Union guidelines for the handling of laboratory animals. All procedures of the protocol were approved by the Committee on the Ethics of Animal Experiments C2EA-26 (n° APAFIS#4617- 2016032118119771 v1) of the Paris-Saclay University and the French Ministry of Research. All efforts were made to minimize animal suffering.

Twelve 6- to 7-week-old conventional C57BL/6 female mice were acquired from Charles River Laboratories (France) and were housed at the animal facility of the Faculty of Pharmacy, Paris-Saclay University (agreement number 92-019-01), with *ad libitum* access to irradiated food and autoclaved water. To induce an intestinal dysbiosis allowing their infection by *C. difficile,* mice received an antibiotic pre-treatment [34] (Figure 1A) (Supplementary methods). Mice were infected by oral gavage with 10^5^ spores each whereas mice from the control group received water. Three mice from the infected groups were euthanized 8h, 28h and 32h post-infection, and the three uninfected mice were euthanized at 8h. Following sacrifice, entire *caeca* were collected and caecal content were sampled to determine the burden of *C. difficile* (Figure 1B). The rest of the *caeca* and their contents were placed in RNA*protect* solution (Qiagen) for further RNA extraction (Supplementary methods).

**Figure 1.**
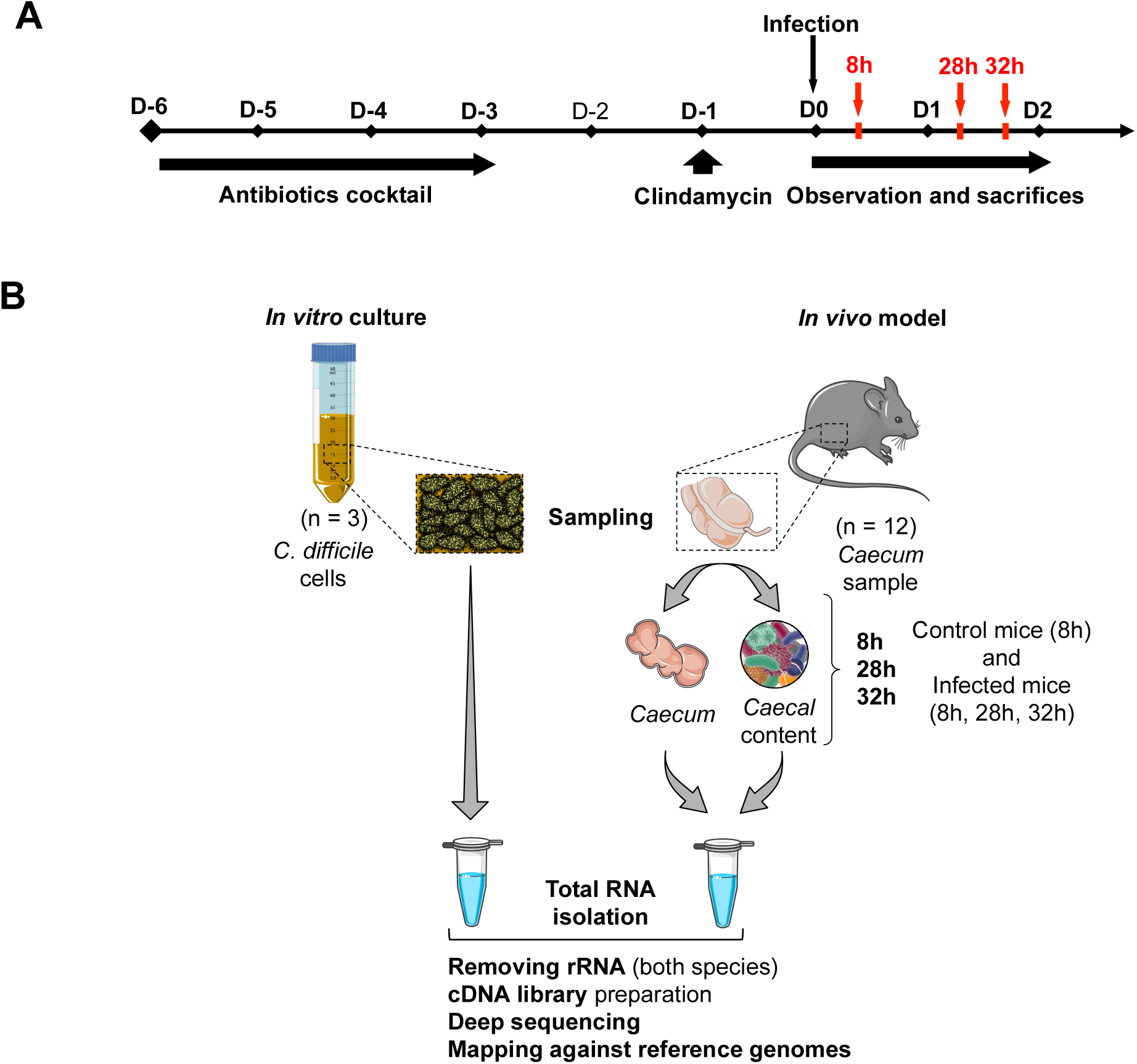
Description of dual RNA-seq experiment workflow. **(A)** The mouse model for *C. difficile* infection used for dual RNA-seq experiment has been described previously by Chen et *al*. [34]. All the mice, including the control mice, received the same antibiotic treatment prior to the infection. Nine mice were infected with a suspension of 10^5^ spores of the *C. difficile* 630Δ*erm* strain and three control mice received sterile water. Three infected mice were euthanized at each time point 8h, 28h and 32h post-infection for RNA extraction. The three uninfected control mice were euthanized 8h after the administration of water. **(B)** All the mice (infected and uninfected) were treated according to the same protocol. For each mouse, the RNAs from the caecum and its content were extracted and purified separately before being assembled in one tube and then sequenced together. RNAs from the *in vitro* culture cells were purified with the same protocol (TRIzol) and sequenced with the same method (Illumina).

### RNA extraction, library preparation and RNA sequencing

For *in vivo* RNA isolation, eukaryotic and prokaryotic cells were lysed separately in Fast-prep apparatus (lysing matrix D and B, respectively, two cycles of 30s at speed 6.5). Total RNA isolation was then performed using Trizol (Sigma) as described previously [35] (Figure 1B). The quality of eukaryotic and prokaryotic total RNA was tested with Agilent RNA 6000 Pico kit and quantified before sequencing. For library preparation, eukaryotic and prokaryotic RNA were mixed in a 2:1 proportion to ensure sufficient genome coverage and maximize the prokaryotic RNA sequencing. Ribosomal RNA depletion and library preparation was done using an Illumina TruSeq Stranded Total RNA kit with 1:1 mixture of human/mouse/rat and bacterial rRNA removal solutions for rRNA depletion. The resulting libraries were multiplexed and sequenced on an Illumina NextSeq 500 system as a paired end (PE) 50-35 nucleotide run using a NextSeq 500/550 High Output 75 cycles v2 reagent kit (Illumina).

### Reads alignment and differential expression analysis

The sequencing data were processed as described in Supplementary methods. Differential expression analysis was performed using the DESeq2 [36] based script SARTools [37], and genes were considered differentially expressed with at least 1 log2 fold change and an adjusted *p*-value < 0.05. All analysis for mouse transcriptome data was performed using R [38] and RStudio [39]. Genes with count mean lower than 10 were discarded for downstream analyses (39,719 to 21,085 genes). TMM normalization was done using edgeR [40] and data were linearized with voom function from limma with quantile normalization [41, 42]. Then we applied a one-way analysis of variance for TREATMENT factor for each gene and made pairwise Tukey’s post hoc tests between groups. Significant gene with *p*-value < 0.05 and fold-change > 2 (or < −2) in at least one comparison were selected for downstream analysis (2,297 genes). For functional enrichment analysis we used MSigDB v7.5 [42] and applied Fisher exact test with false discovery rate (fdr) correction for multiple testing to find significant overlaps.

### Functional classification with MA2HTML

Gene-set enrichment was assessed using functional gene classification from the Ma2HTML database [43] (https://mmonot.eu/MA2HTML/, extraction number 1652263129), and the log2 fold change values between *in vivo* (MI) versus *in vitro* (IV) conditions of each of the 2,855 genes with an associated functional class. For the 20 classes, gene-set enrichment was measured with the blitzGSEA software [44].

### Taxonomic classification of shotgun sequencing reads

Relative bacterial cell abundances were evaluated using mOTUs2 [45]. Briefly, mOTUs2 performs taxonomic classification of shotgun metagenomics and metatranscriptomics sequencing reads using a single-copy, non-16s rRNA marker gene approach. The 10 marker genes used are highly conserved and can serve as good proxies to assess the relative abundances of active cells in the community. Prior to classification, mouse reads were removed of the dataset by two consecutive alignments to the mouse genome (RefSeqs accession GCF_000001635.27) using STAR [46] and Bowtie2 [47]. mOTUs2 was run on each sample read dataset using the following parameters: $ motus profile -s R1_001.fastq -t 10 -k taxonomic_level -B -q -o output_file_ABOND.motus. Intergroup differences at the species level were assessed using the Linear discriminant-analysis effect size (LEfSe) method [48]. LEfSe uses Kruskal-Wallis test (two-tailed nonparametric) to determine the level of significance of differences in features (bacterial taxa) between two conditions.

### Analysis of available *in vivo* transcriptomics data for differential ncRNA expression

For comparative analysis with available *in vivo* raw transcriptomics data from previously reported datasets we chose the conditions that most closely approximated the comparison we realised during the present study: *in vivo* conditions in infected mice vs *in vitro* conditions. We used the *in vivo* WT_RNA 3-days post *C. difficile* infection (European Nucleotide Archive, ENA identifier: PRJNA666929) vs. *in vitro* Base media (ENA identifier: PRJNA667108) data reported by Pruss *et al*. [32] and the days 2 post *C. difficile* infection vs. *in vitro* overnight culture in TY medium (ENA identifier: PRJNA612095) in Fletcher *et al*. [27]. All RNA-seq replicates of each condition were downloaded from ENA (see Table S5) and processed separately for the 2 experiments with a snakemake script [49] on the high-throughput computing resources of the French Institute of Bioinformatics (IFB) (Supplementary methods). Final tests and Figure 5 were produced from differential gene analysis tables using a dedicated R script (with the Eulerr package [50]). The convergence of differential expression experiments was assessed using *χ^2^*tests on differential gene lists. Heatmaps of differential genes in all experiments were drawn (Complex Heatmap R package [51]) in log2 fold change unit, with values set to 0 in case of the absence of differential expression. All codes are available on github [https://github.com/i2bc/Dual_Seq_Cdiff_Mouse].

### NcRNA detection from RNA-seq data using DETR’PROK

The prediction of new RNAs was carried out using shell version 2.1.3 of the DETR’PROK pipeline [52] for two conditions of the three experiments with 630 *C. difficile*: *in vitro* and infected mice samples kIV, kMI from this Kreis *et al* study, *in vitro* Base media (pBase) and *in vivo* WT_RNA 3- days post-infection (pWT) samples from Pruss *et al*. [32], and *in vitro* TY (fwtTY) and 2-days post-infection (fwtd2) samples from Fletcher *et al*. [27] (Supplementary methods and Table S6). DETRPROK predictions for ncRNA for the 6 conditions were combined (clusterize.py, S-MART tools [53]) and candidates overlapping in sense or antisense (CompareOverlapping.py, S-MART tools) with previously annotated rRNAs or tRNAs were removed, yielding 118 potential new transcript candidates.

## Data availability statement

Raw sequencing data have been submitted to ENA with the accession number PRJEB64651.

## Results and Discussion

### Animal model and protocol optimization

Although several *in vivo* transcriptomic studies have already been carried out on *C. difficile* and mice separately [26–29, 32, 54], none have so far looked by RNA-seq in animal model at the expression dynamics of RNAs including ncRNAs simultaneously in the pathogen and the host. We have chosen the conventional mouse model that mimics the conditions of human infection associated with antibiotic pre-treatment in animals [34]. To determine optimum sampling time points after infection to both recover sufficient caecal content for bacterial RNA extraction and to observe the onset of clinical signs for host response, we first set up a clinical follow-up assay on 6 conventional mice that were infected with 630Δ*erm* spores. In this validated model, the clinical signs of CDI appear between 24h and 36h post-infection. After the oral challenge, mice were monitored during 40h to check the occurrence and evolution of the symptoms induced by CDI, the colonisation rate and weight of the infected mice. Animals were observed regularly for signs of disease, including the consistency of their stools and their behaviour, and weighted one a day, to account for normal fluctuations, and a clinical sickness score (CSS) have been established (Supplementary methods and Figure S1). The first symptoms appeared at 32h with soft stools in all mice, reduced activity and hunched posture for a half of the mice. The pic of symptoms was observed at 40h post-infection, with a CSS score of 6-7; all mice had a marked weight loss and caeca showed a haemorrhagic appearance indicating inflammation. The *C. difficile* colonic colonization plateau was reached at 8h post-infection with approximately 10^8^ vegetative forms/g of faeces and starts to decrease with the appearance of diarrhoea leading to the loss of almost all the caecal content at 40h post-infection. We then selected three time points in the infection kinetics for RNA extraction: an early 8h post-infection point associated to the colonization plateau, and two late 28h and a 32h post-infection points where the first symptoms appear associated with an immune response engaged, but when there is still sufficient quantity of caecal content for sampling before excessive diarrhoea.

For dual RNA-seq, three groups of mice (8h, 28h and 32h post-challenge) (Figure 1A), were infected with *C. difficile* spores and the control mice received saline water solution. At each sampling time point, mice were euthanized, and their *caeca* were sampled. Quantification of *C. difficile* caecal burden confirmed the host colonization by an average of 10^8^ CFU/g of faeces of *C. difficile* vegetative cells at the three time points tested (Table S1). We observed diarrhoea and a lack of activity for three infected mice in two different groups: one mouse after 28h of infection and two mice after 32h. These mice also had a smaller, less filled, and inflamed *caecum* compared to the other mice. The Mouse caeca and contents were lysed separately followed by the RNA extraction yielding two RNA samples for each mouse, predominantly eukaryotic or predominantly prokaryotic RNA (averaging 1200 ng/μL and 120 ng/μL, respectively). Samples were mixed for each mouse prior to sequencing in controlled proportions to maximise the amount of prokaryotic RNA sequenced.

### Dual RNA-seq analysis of *C. difficile* infected mice *caecum*

The distribution of reads mapped on reference genomes for each group are illustrated in Figure S2. Total RNAs for host caecal tissue and microbial caecal content containing pathogen were isolated from infected mice at 8h, 28h and 32h post-infection and then analysed by Illumina deep sequencing. The number (*n* = 3) of *C. difficile* and mouse PE reads from 8h, 28h and 32h post-infection mapped on the reference genomes (GCF_000009205.2 and GCF_000001635.27) are indicated in Figure S2A and S2B, respectively. The remaining PE reads were kept for gut microbiota composition analysis (Figure S2C). Finally, total RNAs were also extracted from three independent 8h *in vitro* cultures of *C. difficile* 630Δ*erm* strain and sequenced (Figure S2A). Given the large gap of reads number between the *C. difficile in vitro* and *in vivo* samples, a representative subsampling of 0.5% of *in vitro* reads has been performed to allow their normalization with the *in vivo* samples for further differential gene expression analysis. Due to the low number of reads in the samples extracted at 8h post-infection, only the data obtained at 28h and 32h were retained for differential *C. difficile* gene expression analysis. We decided to combine these samples together for the comparison with the *in vitro* condition since no significant difference between these two data sets was observed. The principal component analysis (PCA) also revealed similarity between these samples, which are 78% separated from the *in vitro* samples by the PC1 axis (Figure S3).

### Microbial community abundance profiling from dual RNA-seq data using mOTUs2

Microbiota composition constitutes a key parameter affecting the development of *C. difficile* inside the gut and individual outcome of infection. Here we took advantage of conventional mouse model to look at the microbiota composition during CDI using dual RNA-seq data. We applied mOTUs2 program well-adapted for taxonomic profiling of microbial community on housekeeping marker genes from transcriptomic data [45]. As for differential analysis of *C. difficile* gene expression, 8h post-infection samples have been excluded from these microbiota analyses. The PCA analysis of bacterial species in mouse gut revealed a cluster of uninfected mice samples clearly separated from infected samples (Figure 2A). Inside the group of infected mice two clusters could be distinguished corresponding to the groups of *C. difficile*-infected mice presenting or not visible symptoms, clearly seen in three-dimensional PCA graph (Figure 2A). On loading PCA plot (Figure S4A), *C. difficile* appears as a discriminant bacterium in infected conditions contributing as expected to the sample separation. Indeed *C. difficile* modulates the composition of microbiota, either directly via production of *p*-cresol [55] or in an indirect way by inducing indole production by *E. coli* [56].

**Figure 2.**
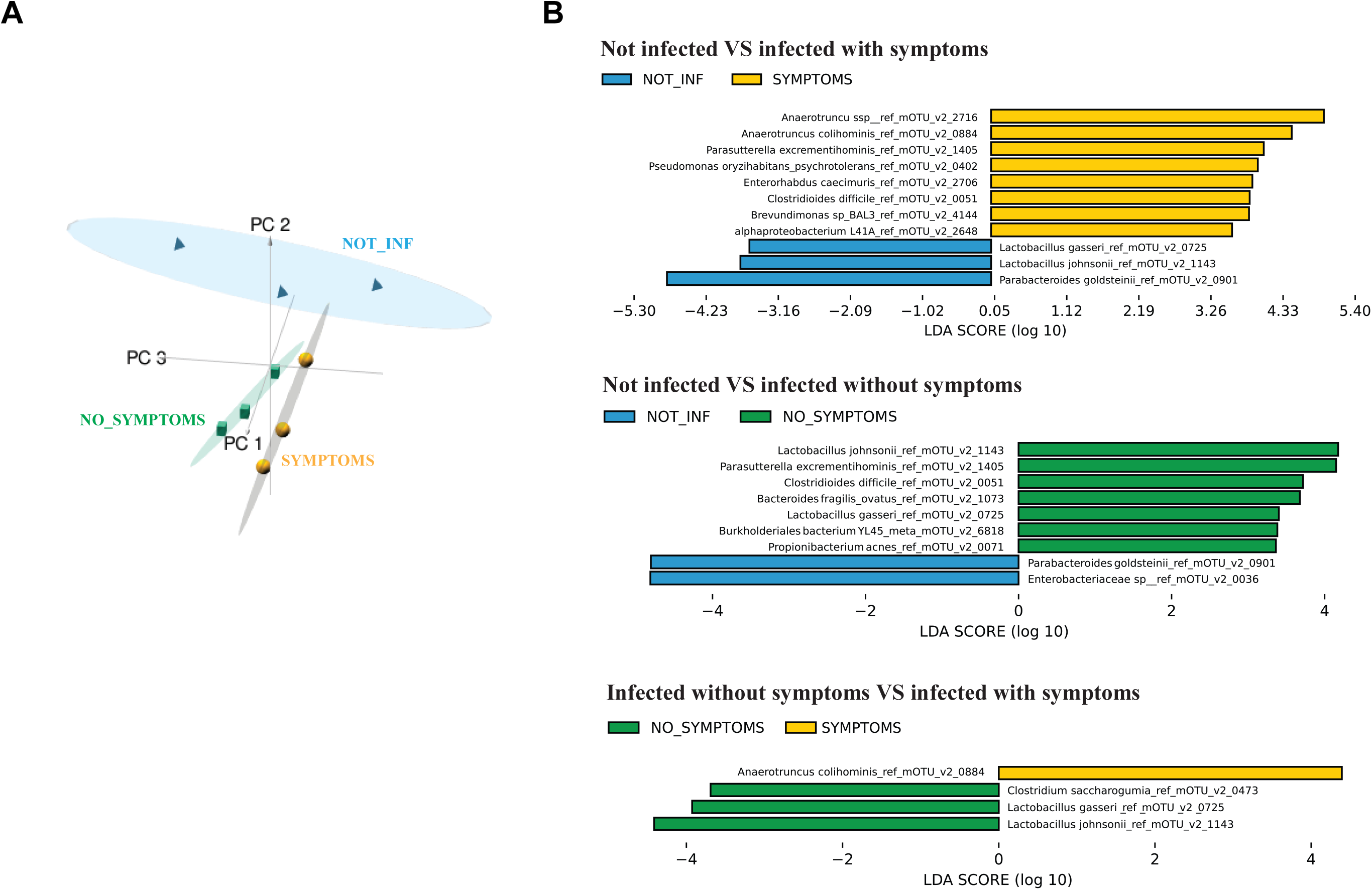
Principal component analysis of the bacterial species identified in the mouse gut (A) and analysis of biomarkers between conditions using LEFSE (B). (**A**) Each symbol on the three dimensional PCA score plot represents a sequenced mouse gut sample. Symbols are colored according to experimental conditions (blue pyramids = NOT_INF: mice not infected with *C. difficile*, yellow spheres = SYMPTOMS: mice infected with *C. difficile* showing visible symptoms, green cubes = NO_SYMPTOMS: mice infected with *C. difficile* showing no visible symptoms). The three main principal components and the corresponding variance proportion are shown. **(B)** Histogram of the LDA scores computed for features differentially abundant between the different conditions being compared. LEfSe score indicates the consistent difference in relative abundance between the features (taxonomic groups) in the microbial communities. The histogram identifies which clades/taxons among all those detected as statistically and biologically differential explain the greatest differences between communities.

The relative community composition at different taxonomic levels is shown in Figure S5. This metatranscriptomic analysis revealed profound alterations in the structure of mouse gut microbiota associated with *C. difficile* colonization and inflammatory symptoms in infected mice, as previously described in other mouse models of CDI [57, 58]. In humans, some common features of dysbiosis have been found in microbiota studies of patients suffering from CDI [57–60]. To search for additional discriminating taxa at species mOTUs level as biomarkers associated with CDI symptoms in our study, we used linear discriminant analysis (LDA) effect size (LEfSe) method [48]. The histograms presented in Figure 2B show the clades that explain the greatest differences between analysed microbial communities. All pairwise comparisons identified as expected *C. difficile* as overabundant species in the infected mice community (Figure 2B). *Lactobacillus* species including *Lactobacillus gasseri* and *Lactobacillus johnsonii* were enriched both in uninfected and symptom-free mice as compared to symptomatic mice and identified as discriminating between non-symptomatic and uninfected samples suggesting potential positive role of *L. gasseri* and *L. johnsonii* in inhibition of CDI symptoms development (Figure 2B and Figure S4B). *Lactobacillus reuteri* was greatly depleted in two symptomatic mice samples 32h post-infection (Figure S4B). Several strains of *L. reuteri* have been previously shown to inhibit *C. difficile* growth *in vitro* [58, 61] but also *in vivo* [62]. In contrast, *L. johnsonii* does not seem to have an effect on the growth of *C. difficile* [62] but its protective effect highlighted in our study could be related to its anti-inflammatory properties [63]. Our analysis also revealed significant association of *Clostridium saccharogumia* with non-symptomatic infected mice as compared to symptomatic mice, and a significant association of *Alistipes indistinctus* with uninfected mice as compared to infected mice (Figures 2B, S4B, and S5A). The *Alistipes* genus was already associated with a protective effect against CDI in a mouse model [64] being an important post-fecal microbiota transplantation (FMT) genus in humans [65].

Relevant to previous observations [27, 66–68], in our study, several *Bacteroidales* species (*Bacteroides dorei/vulgaris* and *B. fragilis*) have been enriched in non-symptomatic infected mice (Figure S4B). In contrast, a group of *Anaerotruncus, Enterorhabdus, Brevundimonas* species was identified as positively associated with *C. difficile* in infected samples, in particular, in symptomatic mice and as depleted in uninfected samples (Figure 2B and Figure S4B). To our knowledge, these genera have not been previously associated, positively or negatively, to *C. difficile*, except for *Anaerotruncus colihominis*. Surprisingly, this Gram-positive rod-shaped bacterium is known to have anti-inflammatory effects, probably by butyric acid production and was long-term increased after FMT in one patient suffering from CDI [69]. Thus, this species deserved more studies to understand the nature of its potential interaction with *C. difficile*.

This microbiota composition profiling allowed clustering the samples into three groups (uninfected mice, symptomatic and non-symptomatic infected mice) to consider during the course of this study exploring the correlation between microbiota structure, *C. difficile* gene expression and mice transcriptomic data.

### Comparative *C. difficile* transcriptomic analysis between *in vitro* culture and infectious conditions

For *C. difficile*, our transcriptomic analysis revealed a total of 1,309 genes (559 upregulated and 750 downregulated (Table S2) exhibiting differential expression between *in vitro* and a 28-32h post-infection condition (Figure 3A) including 61 ncRNA genes (Table 1). All differentially expressed genes were then assigned to functional categories with the MA2HTML database classification (Figure 3B). Gene-set enrichment analysis was performed to compare *C. difficile* gene expression profiles in infected mice and *in vitro* growth conditions. Eleven classes out of 20 have an adjusted *p*-value < 0.05 associated with an fdr < 25% (Table S3). The 2 classes showing the best normalized enrichment score are associated with sporulation as upregulated gene set and regulations as commonly downregulated class (Figure 3B, Figure S6).

**Figure 3.**
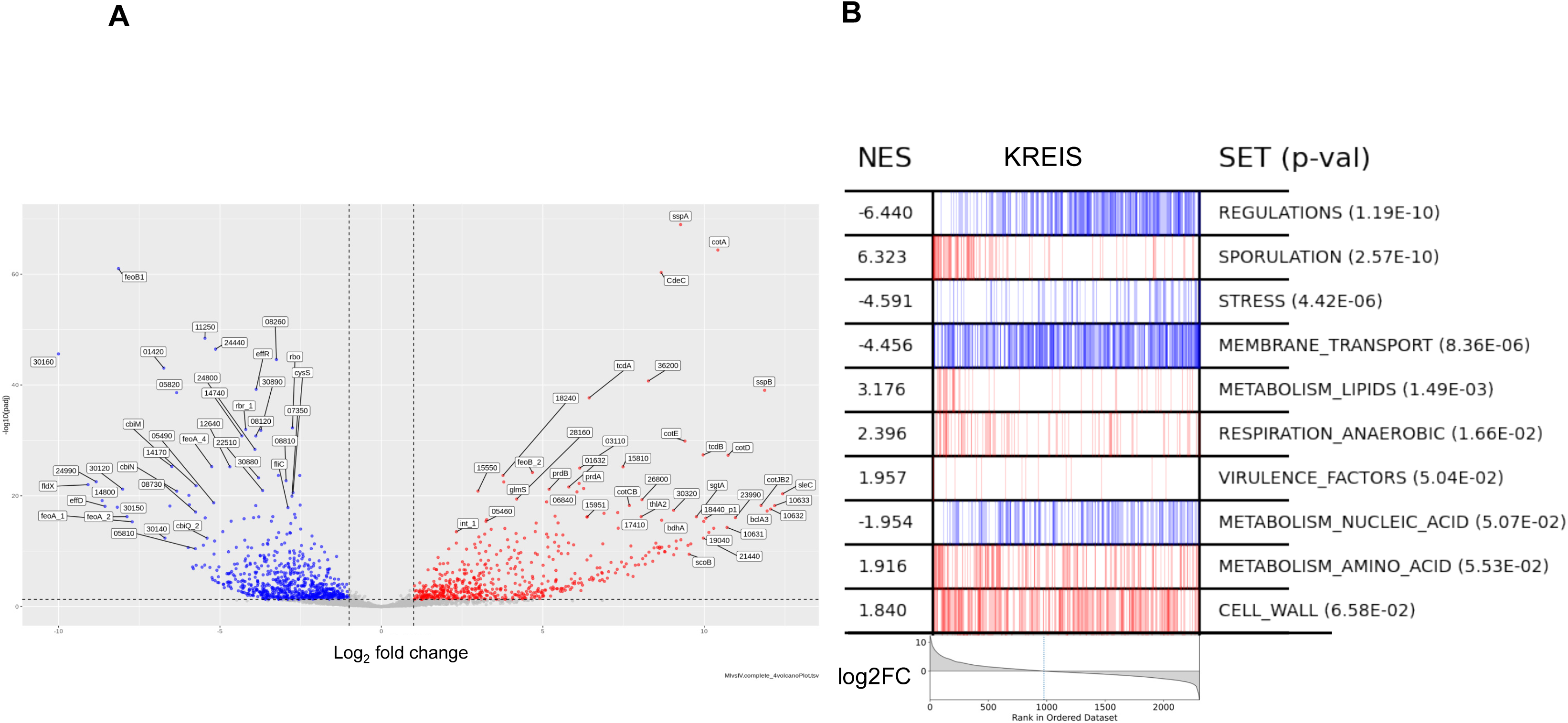
Genes of *C. difficile* differentially expressed between groups of mice infected at 28h and 32h post-infection (MI) and *in vitro* cultures (IV). **(A)** Volcano plot representing the logarithm of the *p*-value adjusted according to the logarithmic ratio (log2 fold change; FC) of the genes differentially expressed between the two conditions. The colored dots correspond to the genes significantly differentially expressed (the genes induced or repressed under the MI conditions in red and blue, respectively). **(B)** Enrichment analysis of Ma2HTML classes with *C. difficile* expression profiles in infected mice versus *in vitro* growth conditions. The enrichment score reflects the concentration on one side of the genes belonging to the class (left side, red for upregulated differentially expressed genes; right side, blue for downregulated differentially expressed genes) as the genes are ordered according to their decreasing log2FC (grey curve at the bottom). NES: normalized enrichment score; SET: class name; *p*-val: adjusted *p*-value.

**Table 1.**
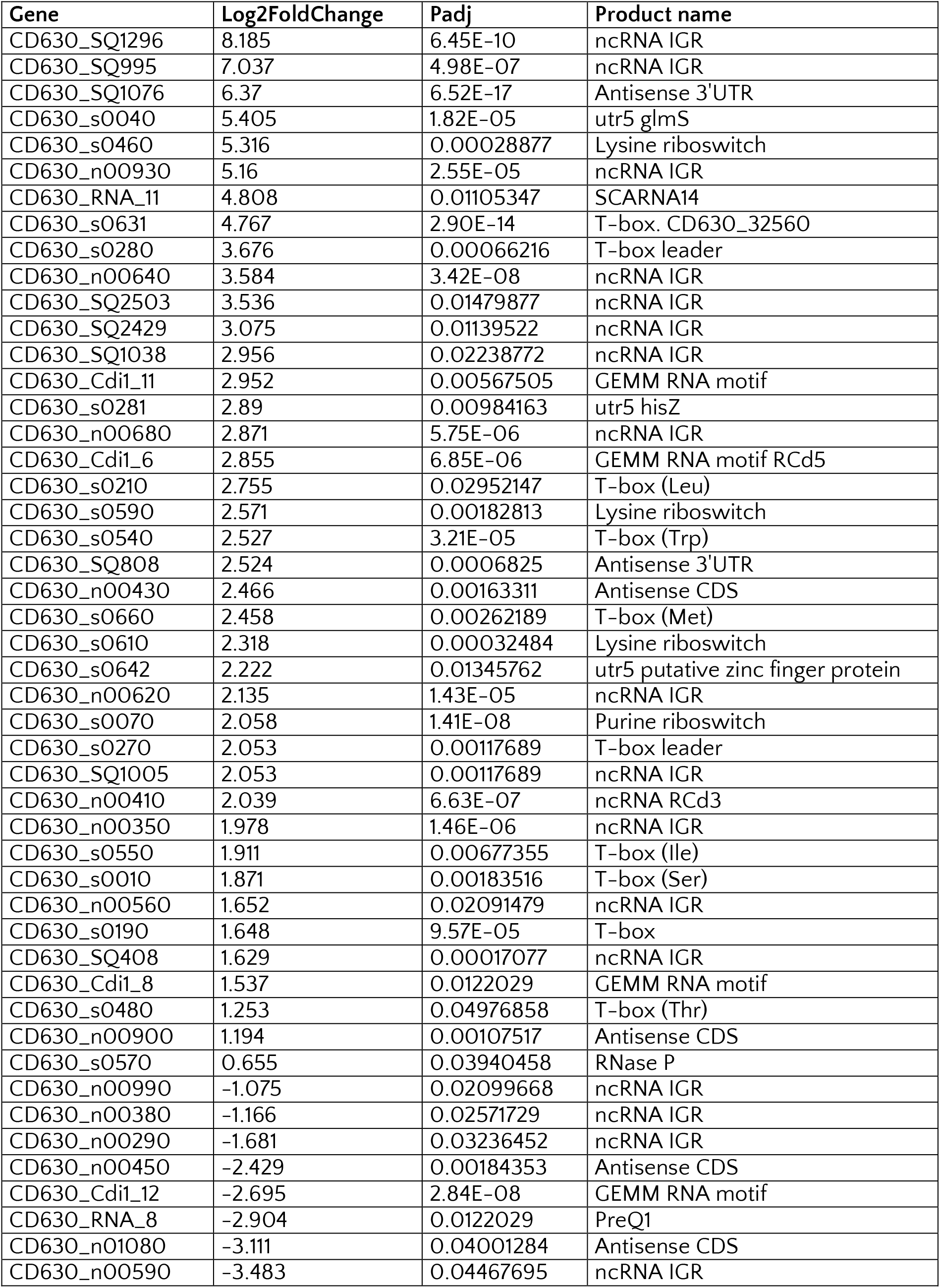
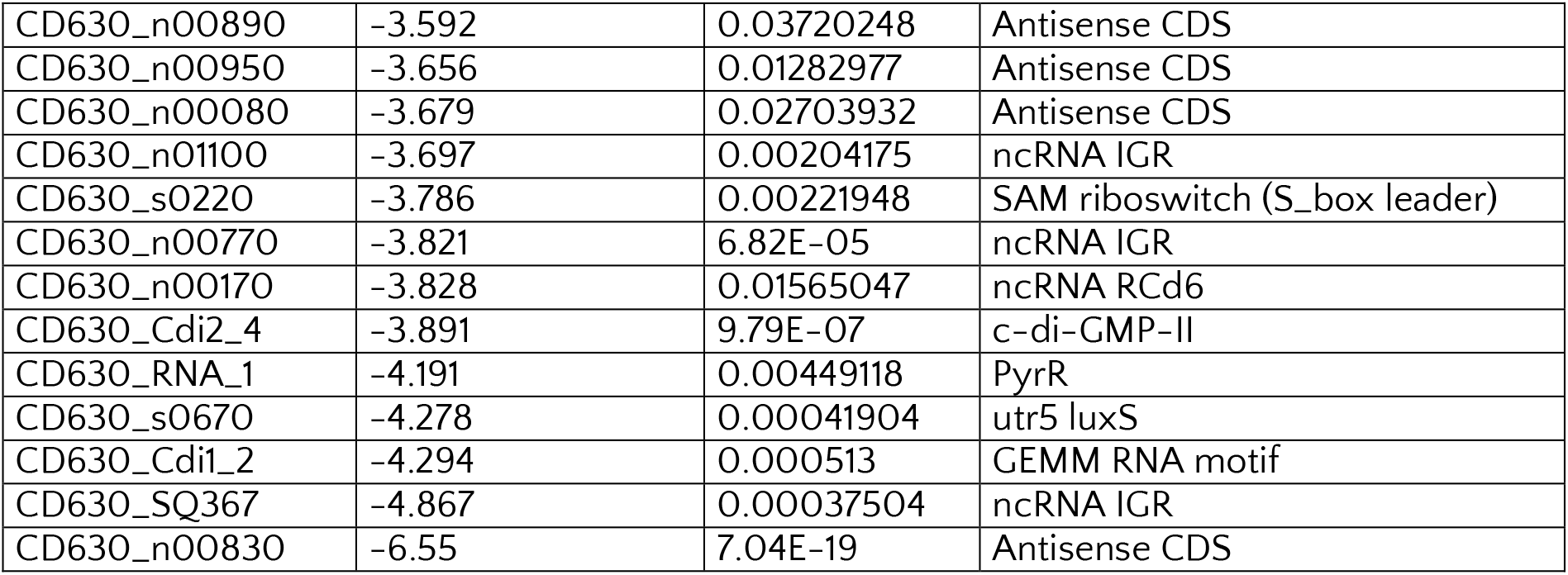
– List of *C. difficile* ncRNAs differentially expressed during infection compared to *in vitro* culture.

Among the genes induced during infection (Table S2) were numerous ribosomal genes reflecting high cellular activity *in vivo* with constant nutrient turnover. The induction of virulence factors such as the toxins TcdA and TcdB (Figure 4A) as well as the adhesin CwpV, promoting self-aggregation and phage resistance of *C. difficile* [70, 71] was in accordance to previous *in vivo* transcriptomics in mice [26, 29, 72] or in pigs [73]. We validated the overexpression of *tcdA* gene in *C. difficile* infected mice as compared to the control by independent qRT-PCR experiment (Figure S7A). In accordance with different expression of flagellar operons in a clinically relevant heat stress [74], some genes from the flagellar assembly F3 operon were induced but several F1 flagellar operon genes were repressed *in vivo* compared to *in vitro* culture. The expression of most of the type IV pilus synthesis genes is decreased *in vivo* in accordance with inverse regulation of flagellum and *pilus* expression by antagonistic type I and type II c-di-GMP-dependent riboswitches [8, 9, 75, 76].

**Figure 4.**
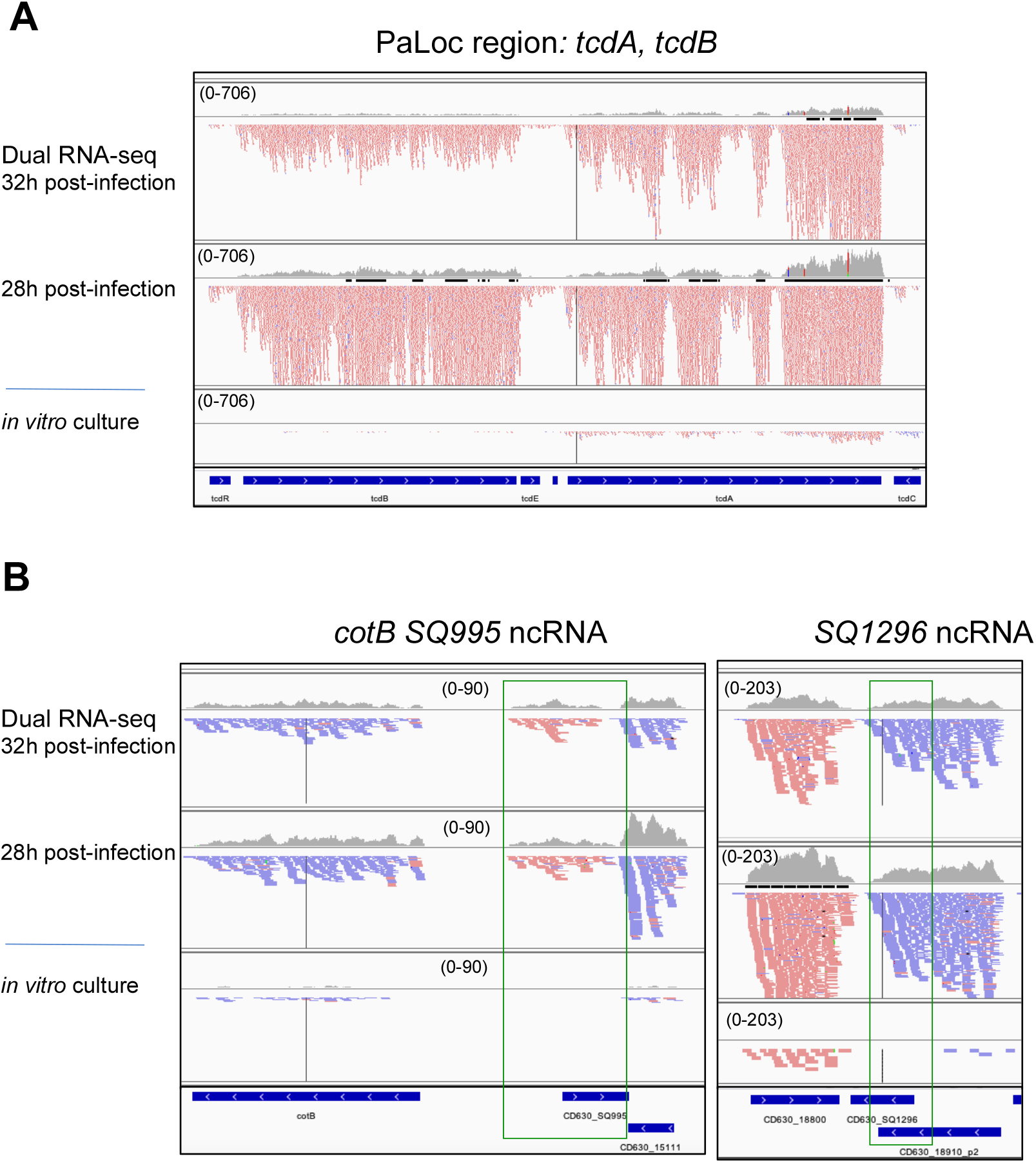
Visualization of dual RNA-seq data for *C. difficile* differentially expressed genes with IGV. Representative examples are presented in **(A)** for protein encoding genes including virulence factors TcdA and TcdB, in **(B)** for ncRNAs. The results for *in vivo* samples 28h post-infection, 32h post-infection are compared with the data from *in vitro* sample. The genomic regions for ncRNAs are presented in a green box. Genes from MaGe annotation are shown at the bottom of each panel. In the IGV visualization profiles, “+” strand reads are shown in red and “-“ strand reads are shown in blue. IGV visualization is presented with adjusted read threshold for each window to compare the data from different samples (scale is indicated as a read threshold range).

**Figure 5.**
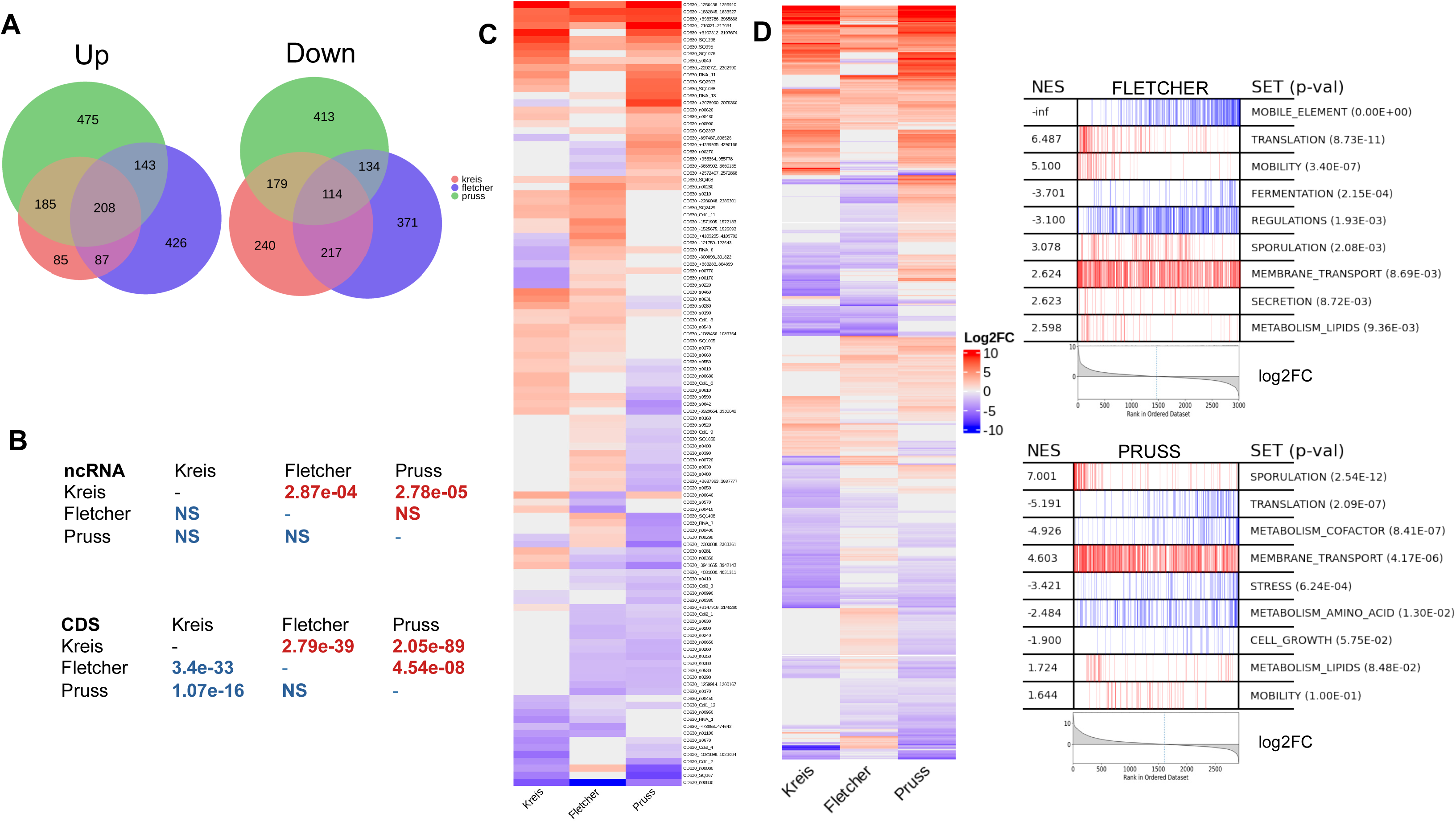
Comparison with available *C. difficile in vivo* transcriptomic data. **(A)** Venn diagram showing the number of differentially expressed genes that are up (left) and down (right) regulated *in vivo* in the infected mice *versus in vitro* growth conditions in this study (in red) as compared to previously reported datasets in Fletcher *et al*. (in blue) [27] and in Pruss *et al*. (in green) [32]. **(B)** *χ^2^* tests of pairwise comparisons of the 3 gene expression experiments in *C. difficile*. The table indicates the *p*-value for ncRNAs or CDS up- and downregulated *in vivo* as compared to *in vitro* conditions (up in top right half in red, down in bottom left half in blue). Experiments were compared pairwise. A low *p*-value indicates a dependency of differential expression up or down status between the experiments. **(C)** Heatmap of the differentially expressed ncRNAs *in vivo* as compared to *in vitro* conditions in present study “Kreis” and previous reports “Fletcher” [27] and “Pruss” [32]. **(D)** Heatmap of the differentially expressed CDS genes *in vivo* as compared to *in vitro* conditions in present study “Kreis” and previous reports “Fletcher” [27] and “Pruss” [32]. On the right is shown the enrichment analysis of Ma2HTML classes with *C. difficile* expression profiles in infected mice versus *in vitro* growth conditions for “Fletcher” [27] and “Pruss” [32] studies to compare with this analysis for present study in Figure 3B. The enrichment score reflects the concentration on one side of the genes belonging to the class (left side, red for upregulated differentially expressed genes; right side, blue for downregulated differentially expressed genes) as the genes are ordered according to their decreasing Log2FC (grey curve at the bottom). NES: normalized enrichment score; SET: class name; *p*-val : adjusted *p*-value.

The adaptive metabolic capabilities of *C. difficile* are a fundamental part of the infectious process [30]. One of the main sources of energy for *C. difficile* comes from the fermentation of carbohydrates and amino acids as an important asset to colonise its niche. Many genes dedicated to carbohydrate transport and metabolism are differentially expressed during infection including the induction of ten genes of PTS (Phosphoenolpyruvate-dependent Phosphotransferase System) for the acquisition and phosphorylation of sorbitol, fructose, mannitol and galactitol, and the repression of about twenty genes for the transport of other sugars such as mannose, lactose, but also glucose and glucosides. These sugars are probably absent in the caecum of mice implying the preferential use of other carbon sources. For example, glucose is totally absorbed in the upper part of the intestine and the dysbiosis induced by antibiotics must considerably limit the degradation of complex fibres contained in the mice’s diet into monosaccharides.

The final stage of glycolysis leads to the production of pyruvate, which is then metabolised by several fermentative pathways for energy production or anabolic reactions. Some of these pathways, which lead to the production of butyrate, ethanol, or butanol, pass through a major intermediary product of bacterial carbon and energy metabolism, acetyl-CoA. Acetyl-CoA is the final degradation product of ethanolamine, an abundant compound *in vivo* that can be derived either from the degradation of cell membranes because of disease or directly from the diet. Remarkably, all genes of the *eut* operon involved in ethanolamine metabolism were strongly overexpressed in mice compared to the *in vitro* condition (Figure S8A). Ethanolamine is a source of carbon and nitrogen for the bacteria. The expression of *eut* operon is repressed by glucose, and the observed overexpression is relevant with the repression of glucose transport systems.

*C. difficile* uses amino acids as an energy source through Stickland reactions for coupled fermentation of two amino acids acting as an electron donor and acceptor, respectively. Several amino acids can be used in the oxidative branch, the only acceptors are glycine and proline. We observed a strong overexpression of both selenoenzyme operons, the proline reductase *prd* (Figure S8A) and the glycine reductase operon. Availability of proline in the host has been shown to modulate the bacterial capacity to infect the mouse [77], and proline and hydroxyproline are major components of collagen that can be released by its degradation to further sustain the growth of *C. difficile* [78].

The acquisition of iron from the environment is vital for most prokaryotes. We observed an overexpression of one of the three *feo* operons (*feo*2) (Figure S8A) involved in the uptake of ferrous iron in many pathogenic bacteria. However, in *C. difficile*, this *feo* operon is neither under the regulation by the iron level nor the global regulator Fur [79, 80]. In *Porphyromonas gingivalis,* an homologous Feo system is involved in manganese import, suggesting that this system could allow the uptake of Mn also in *C. difficile*, with a possible modulation for bacterial virulence since Mn is a cofactor of the toxins A and B. The other genes involved in iron acquisition, notably the ABC transporters capable of transporting ferric iron (*CD630_CD2997- 2999*) are repressed *in vivo*. Our results suggest that *C. difficile* is indeed in an iron-depleted environment in this mouse model of infection in agreement with *in vitro* low iron condition transcriptomics [81], showing an overexpression of the flagellar F3 operon and the concomitant repression of both the flagellar F1 and the pili type IV operons.

As observed in previous *in vivo* transcriptomics [26, 29], many sporulation genes (about 100) were induced *in vivo*, including several σ^K^-dependent genes associated with the synthesis of the outer layers of the spore (cortex, spore coat and exosporium) [82–84] (Figure S8A). These results confirm that sporulation is rapidly induced during infection, allowing *C. difficile* to persist in the host gut and disseminate in the environment, despite the host immune response.

Overall, our results are fully consistent with previous *in vivo* transcriptomic analyses in monoxenic or conventional mice [26, 29, 30], which perfectly validate our model.

### Differential expression of *C. difficile* ncRNAs between *in vitro* culture and infectious conditions

Among the 61 differentially expressed ncRNAs (40 induced and 21 repressed) (Table 1), several have been previously identified in a RNA-Immunoprecipitation sequencing (RIP-seq) experiment as being associated with the Hfq protein [18]. For example, RCd6 is repressed, while CD630_n00930 and CD630_n00620 are induced during infection (Figure S8B). Another Hfq-associated RNA RCd5 upregulated *in vivo* is a type I riboswitch binding to c-di-GMP induced during the stationary phase of growth. Inversely, a type II riboswitch CD630_Cdi2_4 and associated *pilA* gene encoding type IV pilus component are downregulated *in vivo*. Among antisense RNAs with highest differential ratio of expression between *in vitro* and *in vivo* conditions, we identified CD630_SQ1076, a putative antisense RNA of the *map2* gene encoding a methionine aminopeptidase that was upregulated during infection (Figure S8B). The antisense RNA most highly repressed during infection was CD630_n00830, an antisense RNA of the *grdB* gene coding for a subunit of glycine reductase (Figure S8B). Interestingly, the *grdB* gene and associated CD630_n00830 antisense RNA are inversely co-regulated *in vivo* as compared to *in vitro* conditions (Figure S8B). Similar inverse regulation was also observed for the proline reductase operon induced *in vivo* and the antisense RNA overlapping the 3’-end expressed *in vitro* (Figure S8B), consistent with the importance of the use of these amino acids *in vivo* for Stickland reaction. Our analysis revealed several previously uncharacterized ncRNAs as highly upregulated during infection. Among them CD630_SQ995 is located in intergenic region (IGR) between *CD630_1511* and *cotB* gene for a spore outer membrane protein, CotB (Figure 4B) that were also induced *in vivo;* and CD630_SQ1296 is located in IGR between *CD630_1880* gene encoding ketopantoate reductase and *pyrE* gene encoding orotate phosphoribosyltransferase in the vicinity of the sequence coding for a fragment of an ABC transporter (Figure 4B). The overexpression *in vivo* of these two previously uncharacterized ncRNAs has been validated by independent qRT-PCR analysis (Figure S7B and C). CD630_n00640 is also induced *in vivo* and found between conjugative transposon Tn1549-like *CD1878.2* and *CD1878* genes. Altogether, these *in vivo* transcriptomic data represent invaluable resources for further detailed characterization of RNA-based regulatory mechanisms during CDI.

### Comparison with available *C. difficile in vivo* transcriptomic data

Several studies previously explored the *in vivo* transcriptomics of *C. difficile* in mouse model of infection, but the ncRNA genes have not been included into these analyses. We thus selected representative raw RNA-seq datasets from two independent studies [27, 32] for further comparative analysis with the present study (Table S4). This analysis revealed a total of 2,258 and 2,319 *C. difficile* genes differentially expressed *in vivo* as compared to *in vitro* conditions, respectively (1,180 and 1,046 genes upregulated, while 1,078 and 1,273 genes downregulated, respectively). Despite the differences in the experimental conditions and post-infection time points, *χ^2^*tests for pairwise comparisons of the three experiments revealed significant overlap for up- and downregulated genes (Figures 5A, 5B, Table S5) encoding virulence factors, sporulation, stress-response and metabolism-related proteins in accordance with previous reports [26–28, 32]. The functional gene-set enrichment revealed two classes associated with sporulation and lipid metabolism as upregulated in all three studies, while regulations and stress-related genes were downregulated in the present study and in Fletcher *et al*. or in Pruss *et al*. report, respectively (Figures 5D and 3B). Importantly, the analysis of raw sequencing data from three independent studies identified a number of ncRNA genes that were differentially expressed during infection *in vivo* as compared to *in vitro* conditions (Figures 5B, 5C, Table S5). Among them, 38, 54 and 24 were upregulated and 19, 36 and 59 were downregulated in the present study and Fletcher *et al*. and Pruss *et al*. datasets, respectively [27, 32]. Strikingly, pairwise comparison revealed a significant overlap with 22 and 13 upregulated ncRNAs in the present study as compared to Fletcher and Pruss data, respectively (Figure 5B). Importantly, CD630_SQ1296 and CD630_SQ995 ncRNAs expression was highly induced while the expression of CD630_n00830 antisense RNA was highly reduced *in vivo* in all three independent studies (Figure 5C). This comparative analysis strengthens the results of present study identifying several ncRNAs as potential key regulators for *C. difficile* adaptation inside the host.

### Prediction of small transcripts from RNA-seq data

We then took advantage of transcriptomics data from present study combined with raw datasets from Fletcher and Pruss [27, 32] to search for new transcripts in *C. difficile* using DETR’PROK pipeline [52]. By combining the three independent datasets for *in vivo* and *in vitro* conditions, this analysis revealed 118 potential new transcript candidates in *C. difficile* including 12 transcripts overlapping annotated CDS and 106 potential new ncRNA genes. Among them 83 potential ncRNAs were predicted in antisense orientation to annotated genes and 23 without overlap with annotated genes (Table S6). Fifty-six of new ncRNAs were identified in antisense orientation to CDS, while 20 corresponded to antisense RNAs for previously identified ncRNAs and 7 were identified in antisense orientation to both CDS and ncRNAs. A number of these potential antisense RNAs have been detected in our Hfq RIP-seq analysis [18]. Among them, an antisense RNA for *CD3236 prdF* proline reductase gene was repressed during infection (Figure S8B). Interestingly, 12 antisense transcripts have been detected for riboswitches, 3 for CRISPR RNAs and 6 for sRNAs associated with Hfq RIP-seq signal. Three antisense RNAs corresponded to previously identified type I toxin-antitoxin system components [15, 16, 85] missing from current NCBI annotation. Eleven predicted RNAs corresponded to longer transcripts overlapping previously annotated ncRNAs or CDS with 3’UTR. Interestingly, in accordance with Hfq RIP-seq analysis [18] DETR’PROK pipeline detected 3 new ncRNAs in IGR of *CD2217*- *CD2218, CD2610* and *CD3364-CD3365* genes and 12 additional IGR ncRNAs associated with lower signal. Altogether these analyses contributed to the definition of the *C. difficile* transcriptomics landscape highlighting the extent of antisense transcription, specifying the boundaries of some previously annotated transcripts and enriching the *C. difficile* ncRNA repertoire for future studies.

### Transcriptomic analysis of host response to *C. difficile* infection

On the mouse side, our analysis revealed 2,297 genes significantly differentially expressed (fold change < −2 or > 2, *p*<0.05) between all conditions. Among them, 800 correspond to regulatory RNA genes, 294 induced and 506 repressed, with mostly lncRNAs (788 differentially expressed, 293 induced and 495 repressed) and few microRNAs (12 differentially expressed, 11 induced and one repressed). The heatmap summarizes the comparison of each mouse transcriptome (Figure 6) and shows the expression profile of the uninfected (red), 8h (blue), 28h (green) and 32h (orange) post-infection mice. Depending on their differential expression, these 2,297 genes could be grouped in 12 clusters, numbered I to XII (Figure 6). Following this overview, we focused on main transcriptomic differences between groups.

**Figure 6.**
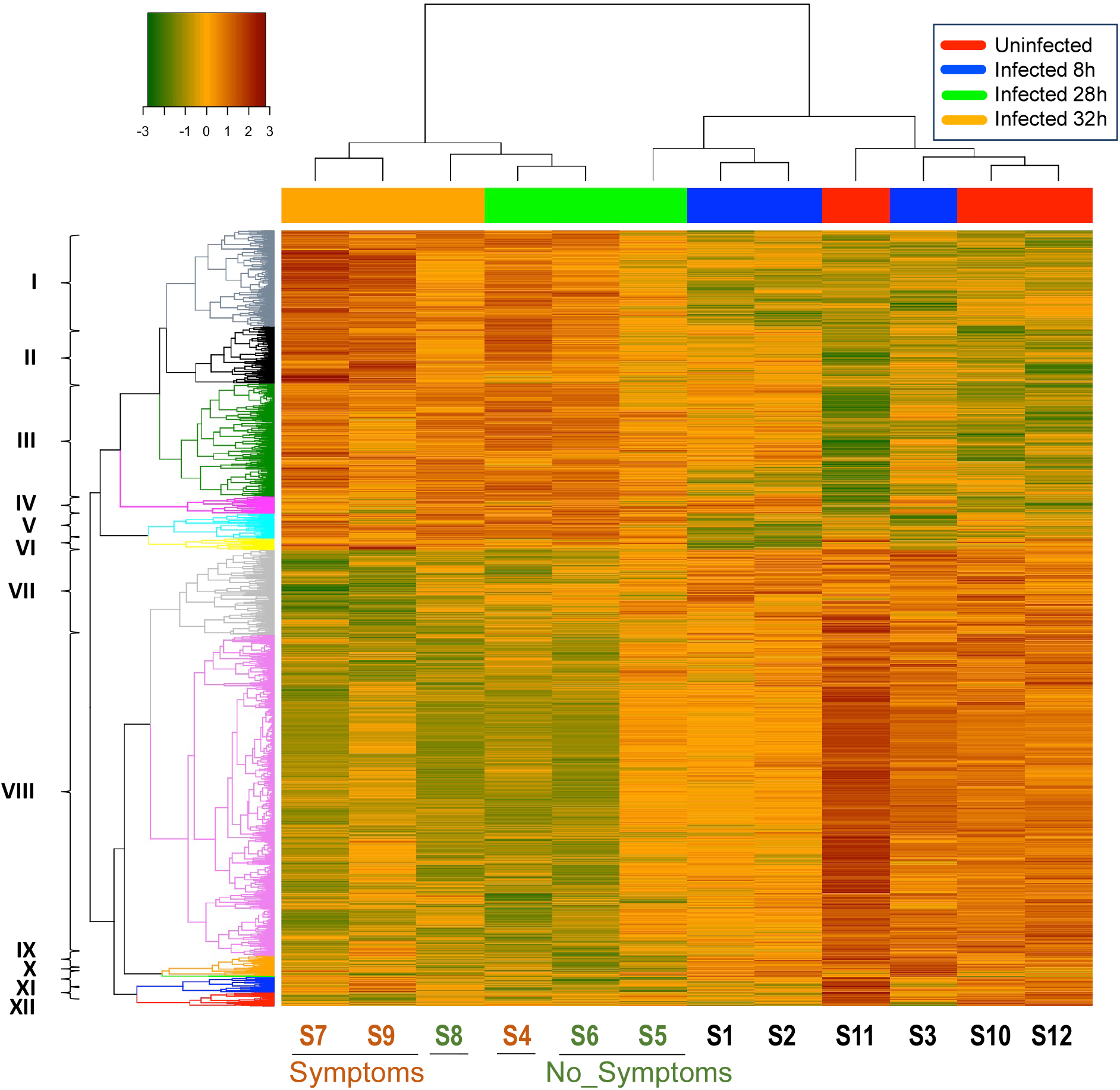
Differential Gene Analysis of mouse genes expression during *C. difficile* infection. An hierarchically clustered heatmap and dendrogram show the expression patterns of the genes differentially expressed in mice between uninfected control mice (Uninfected, samples S11, S12, S13), infected mice at 8h (Infected 8h, samples S1, S2, S3), 28h (Infected 28h, samples S4, S5, S6) and 32h (Infected 32h, samples S7, S8, S9) post- infection. Symptoms: mice infected with *C. difficile* showing visible symptoms (samples S4, S7, S9). No_Symptoms: mice infected with *C. difficile* showing no visible symptoms (samples S5, S6, S8). Clusters of genes are numbered from I to XII and discussed in the text. The color key represents the level of expression for each gene.

For each gene cluster, enrichment analyses identified several differentially expressed pathways or gene families discriminating infected and uninfected mice. As expected, several specific host inflammatory markers were induced during CDI, including members of TNFα signalling pathway (IL-1β, NLRP3, TNF, CCRL2, NFKBIA and FOS), and chemokines (CCL4 and CCXL1) (Table S7). These inflammatory markers from clusters I and II (Figure 6), were particularly induced in 28h and 32h infected mice with clinical signs (diarrhoea) and a highly inflamed cecum, unlike the other symptom-free 28h and 32h infected mice. The strong overexpression of several inflammatory markers in these sick mice revealed a stronger immune response to CDI consistent with the strong caecal inflammation visually observed during animal sacrifice. This expression profile, with the induction of TNFα, IL-1β or CCL4, reveals the activation of a T Helper type 1 (Th1) immune response, while no evidence of Th2 response was observed.

Gene clusters III, V and VI were also induced in the 28h and 32h infected mice both symptomatic and symptom-free. Genes from cluster III are involved in cell division and DNA repair including Nupr1, involved in regulation of cellular catabolic process or programmed cell death, already shown as part of the host response to bacterial infection [86, 87].

Few genes (cluster IV) show a distinct profile, with an induction in all infected (8h, 28h and 32h) mice. Most of these genes are part of metabolism pathways, but two could also be related to immune response including BOLA2 (Bovine leukocyte antigen family member 2), upregulated in CD4+ T cells by JAK-STAT signalling following IL-12 stimulation, and then Th1 immune response [88], and GNAT3 (G Protein Subunit Alpha Transducin 3) encoding a taste receptor, which is also expressed in the gut with potential role in innate immunity [89].

Among genes repressed during CDI (clusters VII – XII), we found many genes encoding proteins involved in: i) metabolism (fatty acid metabolism, cholesterol homeostasis, glycan and glycosaminoglycan metabolism, …); ii) cell junction interactions and cell adhesion molecule (cadherin, claudin, contactin, …); iii) signal transduction (ligand-gated ion channel as glutamate receptor, adrenergic receptor and calcium channel) (Table S8). Similar results were obtained in dual RNA-seq experiments with *Yersinia pseudotuberculosis* proliferating in the gut-associated lymphoid tissue or *Eimeria tenella* infected caecal tissue with shutdown of pivotal cellular functions in response to the infection [23, 90].

As no statistically significant differences in gene expression were observed with Principal Component Analysis (Figure S9A) between 28h and 32h infected mice, these two groups could be combined into a late infected mice group. 1,780 genes were significantly differentially expressed between late infected mice and uninfected mice (Figures S9A and S10A) (530 repressed and 1,250 induced during infection). Among the most induced genes in infected mice were host immune response genes encoding TIRAP (TIR adaptor protein), FOS (transcription factor), NLRP3 (member of the NLRP3 inflammasome complex) and several cytokines (TNFα, IL-1α, IL-1β, IL-6, IL-22, CXCL1-2-5, CCL2-3-4-7). We validated the overexpression of IL-1β, Il- 22, CXCL5 gene in late infected mice as compared to uninfected mice by independent qRT-PCR experiment. Several genes encoding anti-microbial peptides, such as α-defensins or intelectin-1 were repressed in late infected mice [91]. It has been previously observed that a parasite, *Cryptosporidium parvum*, was able to downregulate these genes as another immune evasion strategy [92].

The large number of reads aligning to the mouse genome allows for more detailed comparisons between different groups of samples. We therefore looked for potential differences based on kinetic or clinical criteria (Figure S10B and S10C), by comparing the expression profile of late infected (28h and 32h) vs. early (8h) infected mice and of symptomatic (sick) vs. asymptomatic (healthy) mice within the late infected mice group.

When the combined group of 28h and 32h late-infected mice was compared with the 8h early infected mice group (Figure S9B), we found a much lower number of differentially expressed genes than for the previous analysis (late infected mice versus uninfected mice). Only 375 genes were significantly differentially expressed (98 repressed and 277 induced) suggesting the quick induction of host response to CDI with the majority of genes differentially expressed as early as 8h post infection (e.g. inflammatory response *TNFα, IL-6* and *IL-22*), although the histological and clinical consequences of the inflammatory process are not yet observable.

Finally, within the late infected mice (28h and 32h), we were able to compare gene expression between sick mice vs. symptom-free mice. We identified as many as 3,538 significantly differentially expressed genes in the sick animals, 1,909 induced and 1,629 repressed (Figure S9C). The differentially expressed genes were classified into biological functions using Reactome [93]. A large number of immune and inflammatory responses genes were induced in symptomatic mice compared to asymptomatic mice. Other induced genes are involved mostly in metabolism, homeostasis, signal transduction, keratinization and tissue remodelling (Table 2). Several genes encoding metalloproteases with collagenase activity, notably the *MMP8* gene, were strongly overexpressed in sick mice, in accordance with the results of Fletcher *et al*. [27]. Degradation of collagen by these host proteins may participate to the tissue lesions but may also sustain the growth of *C. difficile* by providing proline and hydroxyproline to the bacteria.

**Table 2.**
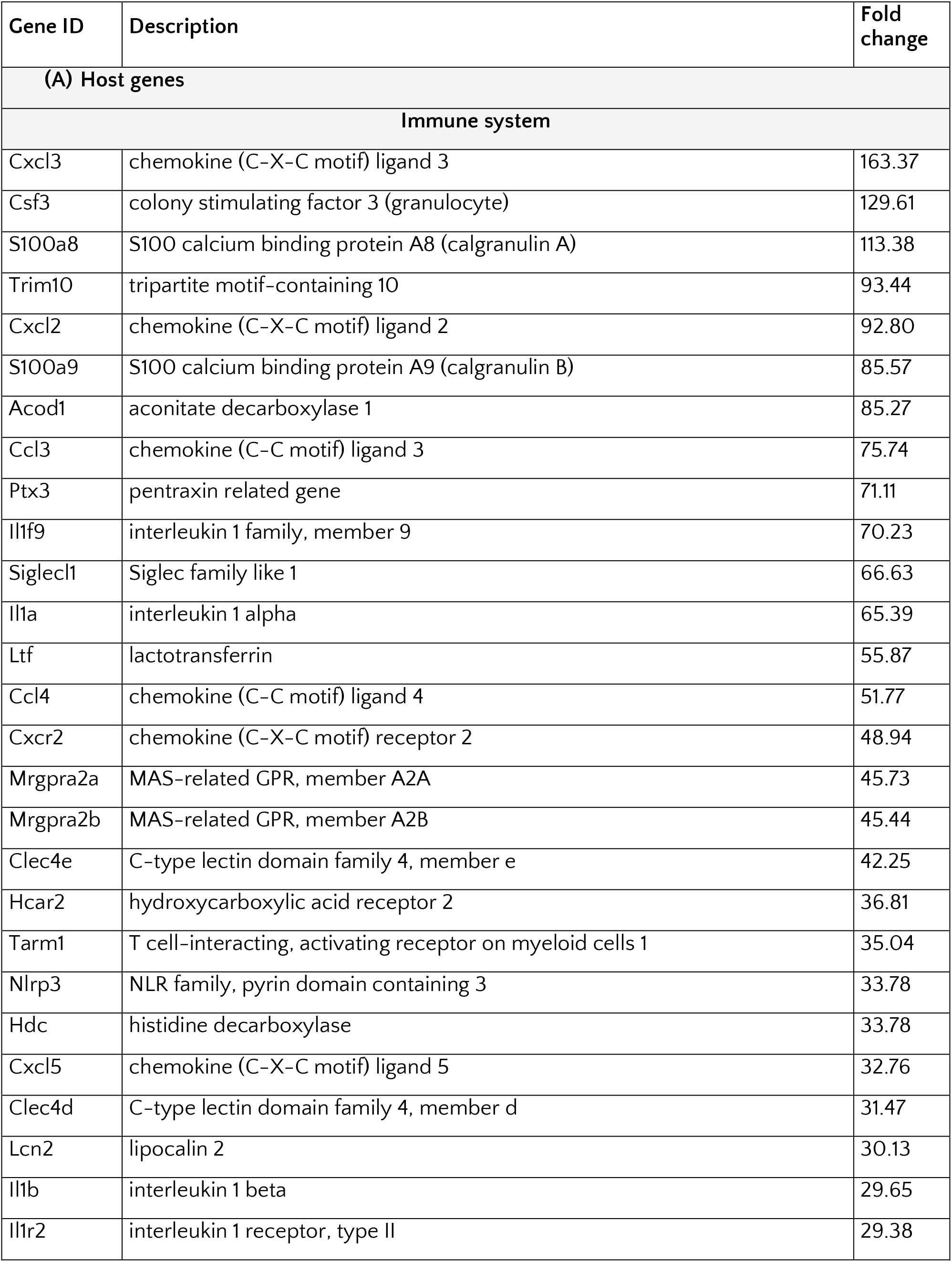
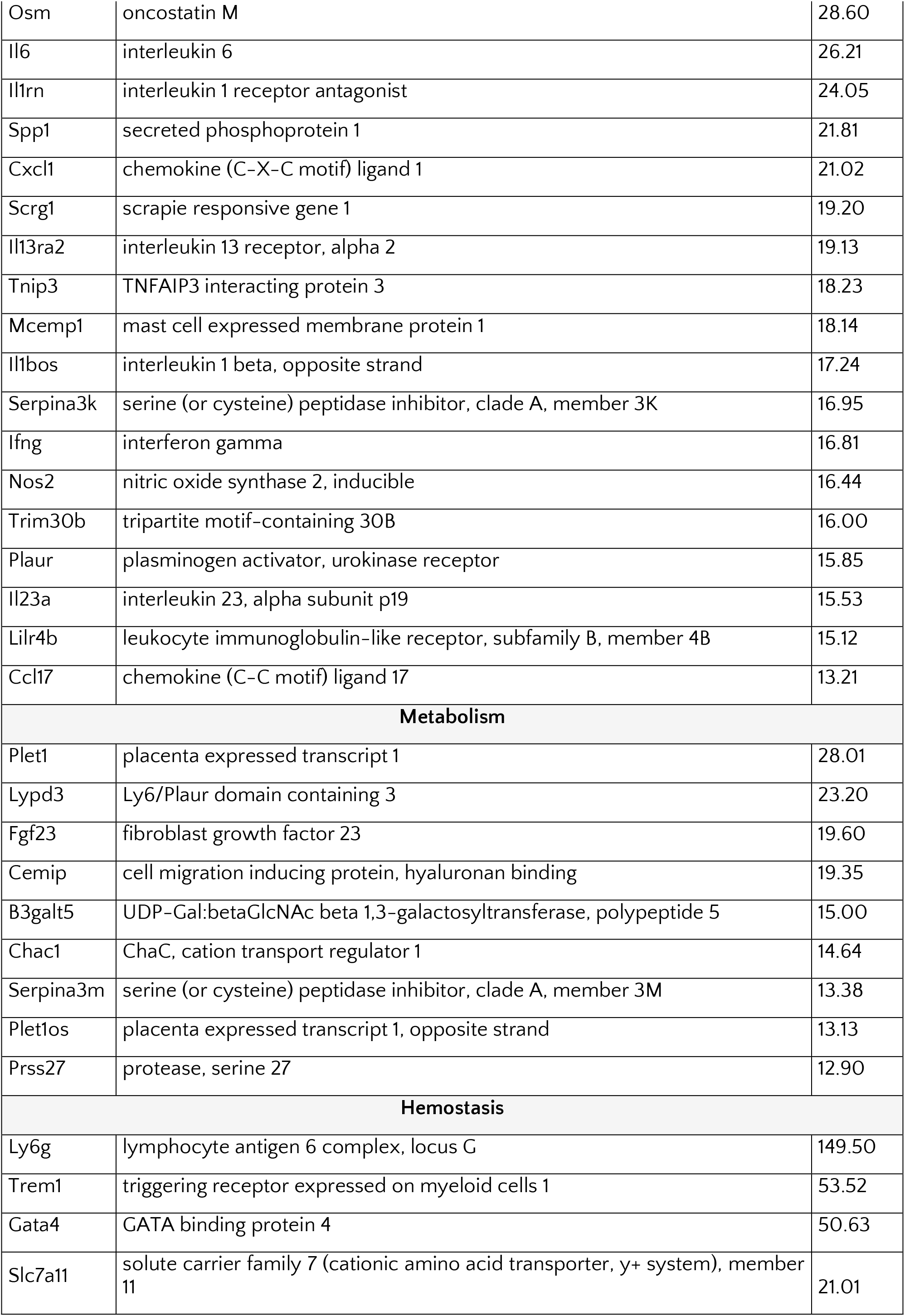
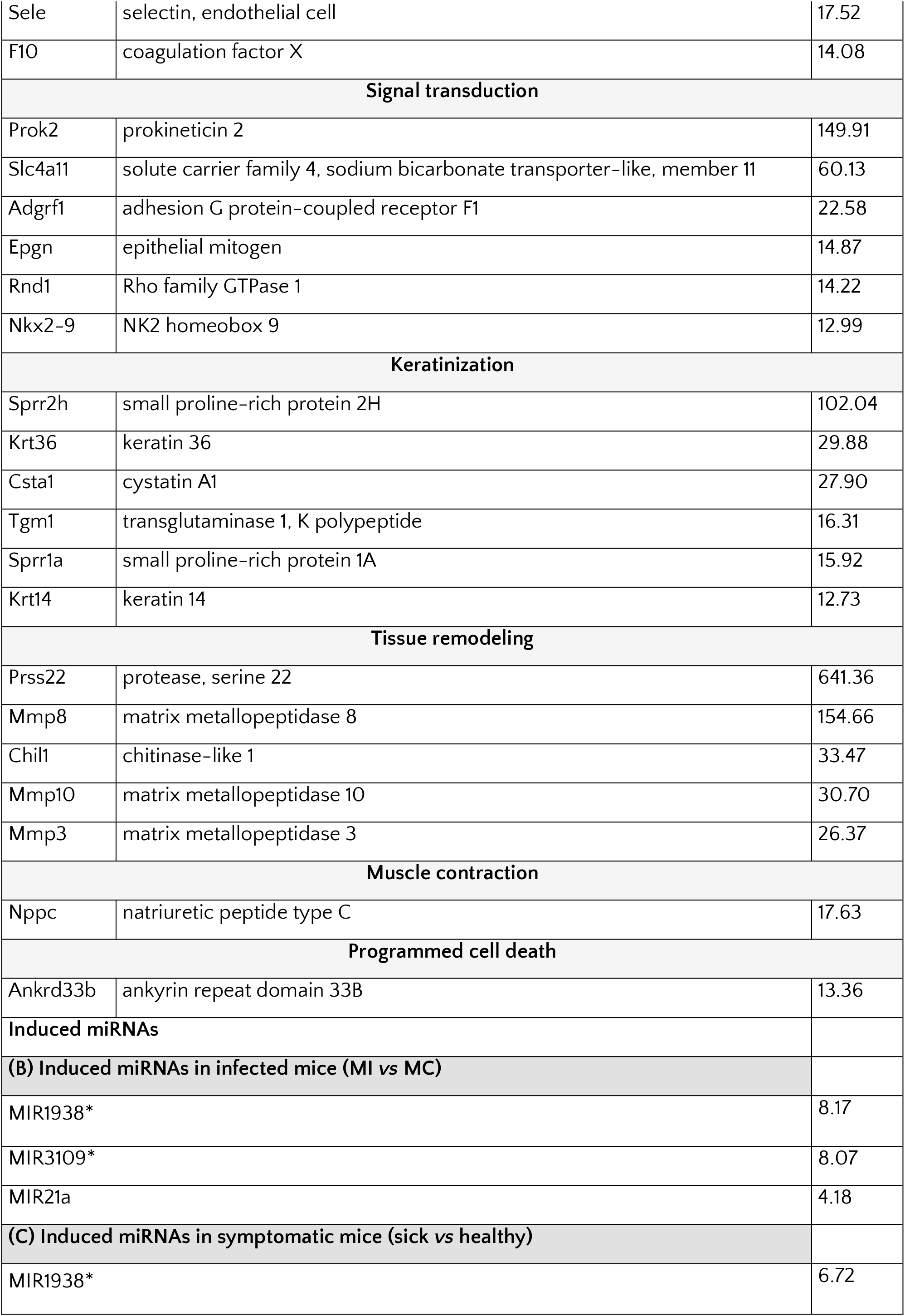
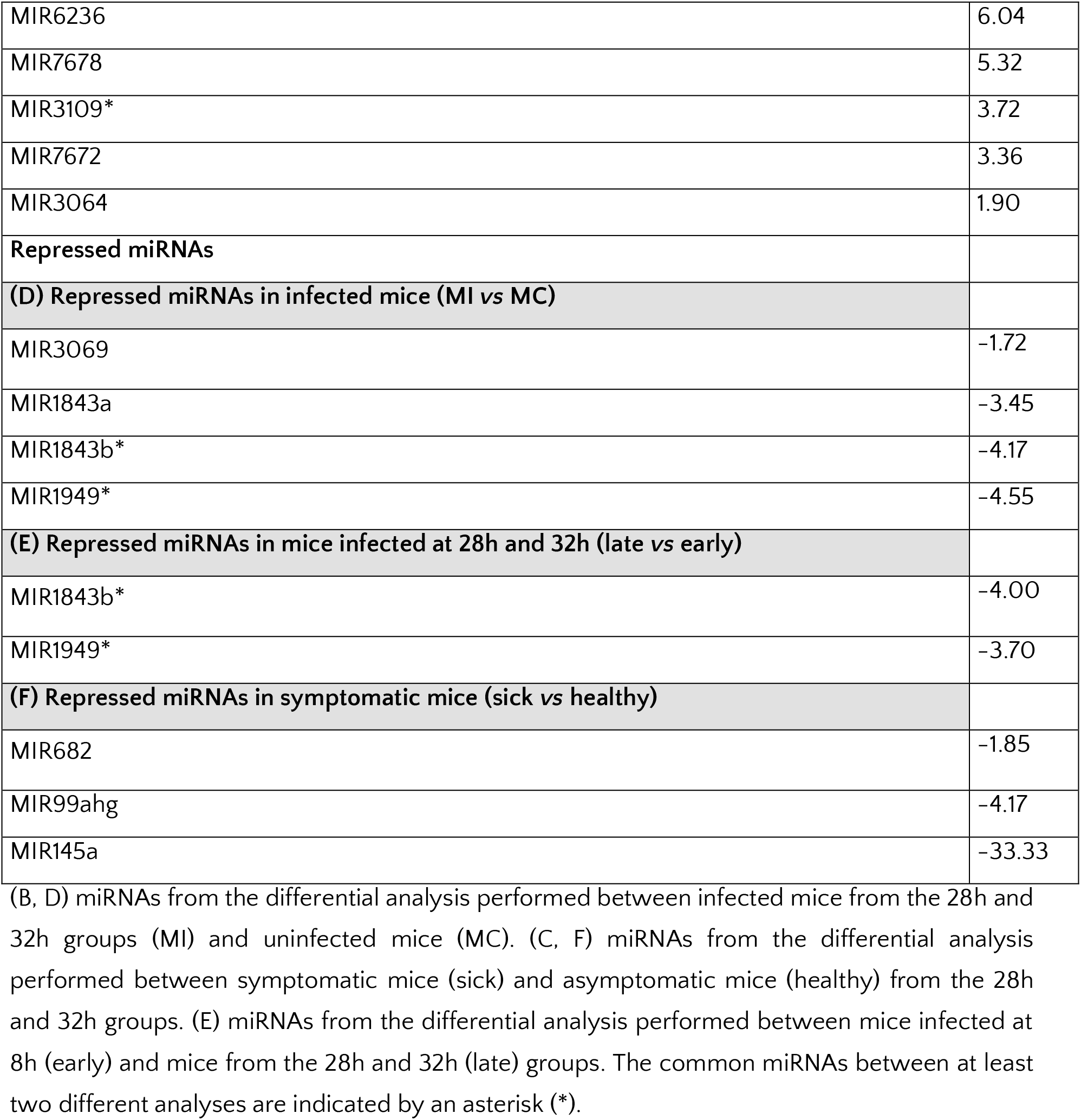
Host genes strongly induced in sick symptomatic mice, compared to asymptomatic mice (both being late infected animals) (A) and miRNAs induced (B, C) or repressed (D, E, F) in mice during CDI.

| Gene ID | Description | Fold change |
| --- | --- | --- |
| **(A) Host genes** |  |  |
| **Immune system** |  |  |
| Cxcl3 | chemokine (C-X-C motif) ligand 3 | 163.37 |
| Csf3 | colony stimulating factor 3 (granulocyte) | 129.61 |
| S100a8 | S100 calcium binding protein A8 (calgranulin A) | 113.38 |
| Trim10 | tripartite motif-containing 10 | 93.44 |
| Cxcl2 | chemokine (C-X-C motif) ligand 2 | 92.80 |
| S100a9 | S100 calcium binding protein A9 (calgranulin B) | 85.57 |
| Acod1 | aconitate decarboxylase 1 | 85.27 |
| Ccl3 | chemokine (C-C motif) ligand 3 | 75.74 |
| Ptx3 | pentraxin related gene | 71.11 |
| Il1f9 | interleukin 1 family, member 9 | 70.23 |
| Siglecl1 | Siglec family like 1 | 66.63 |
| Il1a | interleukin 1 alpha | 65.39 |
| Ltf | lactotransferrin | 55.87 |
| Ccl4 | chemokine (C-C motif) ligand 4 | 51.77 |
| Cxcr2 | chemokine (C-X-C motif) receptor 2 | 48.94 |
| Mrgpra2a | MAS-related GPR, member A2A | 45.73 |
| Mrgpra2b | MAS-related GPR, member A2B | 45.44 |
| Clec4e | C-type lectin domain family 4, member e | 42.25 |
| Hcar2 | hydroxycarboxylic acid receptor 2 | 36.81 |
| Tarm1 | T cell-interacting, activating receptor on myeloid cells 1 | 35.04 |
| Nlrp3 | NLR family, pyrin domain containing 3 | 33.78 |
| Hdc | histidine decarboxylase | 33.78 |
| Cxcl5 | chemokine (C-X-C motif) ligand 5 | 32.76 |
| Clec4d | C-type lectin domain family 4, member d | 31.47 |
| Lcn2 | lipocalin 2 | 30.13 |
| Il1b | interleukin 1 beta | 29.65 |
| Il1r2 | interleukin 1 receptor, type II | 29.38 |
| Osm | oncostatin M | 28.60 |
| Il6 | interleukin 6 | 26.21 |
| Il1rn | interleukin 1 receptor antagonist | 24.05 |
| Spp1 | secreted phosphoprotein 1 | 21.81 |
| Cxcl1 | chemokine (C-X-C motif) ligand 1 | 21.02 |
| Scrg1 | scrapie responsive gene 1 | 19.20 |
| Il13ra2 | interleukin 13 receptor, alpha 2 | 19.13 |
| Tnip3 | TNFAIP3 interacting protein 3 | 18.23 |
| Mcemp1 | mast cell expressed membrane protein 1 | 18.14 |
| Il1bos | interleukin 1 beta, opposite strand | 17.24 |
| Serpina3k | serine (or cysteine) peptidase inhibitor, clade A, member 3K | 16.95 |
| Ifng | interferon gamma | 16.81 |
| Nos2 | nitric oxide synthase 2, inducible | 16.44 |
| Trim30b | tripartite motif-containing 30B | 16.00 |
| Plaur | plasminogen activator, urokinase receptor | 15.85 |
| Il23a | interleukin 23, alpha subunit p19 | 15.53 |
| Lilr4b | leukocyte immunoglobulin-like receptor, subfamily B, member 4B | 15.12 |
| Ccl17 | chemokine (C-C motif) ligand 17 | 13.21 |
| **Metabolism** |  |  |
| Plet1 | placenta expressed transcript 1 | 28.01 |
| Lypd3 | Ly6/Plaur domain containing 3 | 23.20 |
| Fgf23 | fibroblast growth factor 23 | 19.60 |
| Cemip | cell migration inducing protein, hyaluronan binding | 19.35 |
| B3galt5 | UDP-Gal:betaGlcNAc beta 1,3-galactosyltransferase, polypeptide 5 | 15.00 |
| Chac1 | ChaC, cation transport regulator 1 | 14.64 |
| Serpina3m | serine (or cysteine) peptidase inhibitor, clade A, member 3M | 13.38 |
| Plet1os | placenta expressed transcript 1, opposite strand | 13.13 |
| Prss27 | protease, serine 27 | 12.90 |
| **Hemostasis** |  |  |
| Ly6g | lymphocyte antigen 6 complex, locus G | 149.50 |
| Trem1 | triggering receptor expressed on myeloid cells 1 | 53.52 |
| Gata4 | GATA binding protein 4 | 50.63 |
| Slc7a11 | solute carrier family 7 (cationic amino acid transporter, y+ system), member 11 | 21.01 |
| Sele | selectin, endothelial cell | 17.52 |
| F10 | coagulation factor X | 14.08 |
| **Signal transduction** |  |  |
| Prok2 | prokineticin 2 | 149.91 |
| Slc4a11 | solute carrier family 4, sodium bicarbonate transporter-like, member 11 | 60.13 |
| Adgrf1 | adhesion G protein-coupled receptor F1 | 22.58 |
| Epgn | epithelial mitogen | 14.87 |
| Rnd1 | Rho family GTPase 1 | 14.22 |
| Nkx2-9 | NK2 homeobox 9 | 12.99 |
| **Keratinization** |  |  |
| Spr2h | small proline-rich protein 2H | 102.04 |
| Krt36 | keratin 36 | 29.88 |
| Csta1 | cystatin A1 | 27.90 |
| Tgm1 | transglutaminase 1, K polypeptide | 16.31 |
| Spr1a | small proline-rich protein 1A | 15.92 |
| Krt14 | keratin 14 | 12.73 |
| **Tissue remodeling** |  |  |
| Prss22 | protease, serine 22 | 641.36 |
| Mmp8 | matrix metalloproteinase 8 | 154.66 |
| Chil1 | chitinase-like 1 | 33.47 |
| Mmp10 | matrix metalloproteinase 10 | 30.70 |
| Mmp3 | matrix metalloproteinase 3 | 26.37 |
| **Muscle contraction** |  |  |
| Nppc | natriuretic peptide type C | 17.63 |
| **Programmed cell death** |  |  |
| Ankrd33b | ankyrin repeat domain 33B | 13.36 |
| **Induced miRNAs** |  |  |
| **(B) Induced miRNAs in infected mice (MI vs MC)** |  |  |
| MIR1938* |  | 8.17 |
| MIR3109* |  | 8.07 |
| MIR21a |  | 4.18 |
| **(C) Induced miRNAs in symptomatic mice (sick vs healthy)** |  |  |
| MIR1938* |  | 6.72 |

|  |  |
| --- | --- |
| MIR6236 | 6.04 |
| MIR7678 | 5.32 |
| MIR3109* | 3.72 |
| MIR7672 | 3.36 |
| MIR3064 | 1.90 |
| **Repressed miRNAs** |  |
| **(D) Repressed miRNAs in infected mice (MI vs MC)** |  |
| MIR3069 | -1.72 |
| MIR1843a | -3.45 |
| MIR1843b* | -4.17 |
| MIR1949* | -4.55 |
| **(E) Repressed miRNAs in mice infected at 28h and 32h (late vs early)** |  |
| MIR1843b* | -4.00 |
| MIR1949* | -3.70 |
| **(F) Repressed miRNAs in symptomatic mice (sick vs healthy)** |  |
| MIR682 | -1.85 |
| MIR99ahg | -4.17 |
| MIR145a | -33.33 |

(B, D) miRNAs from the differential analysis performed between infected mice from the 28h and 32h groups (MI) and uninfected mice (MC). (C, F) miRNAs from the differential analysis performed between symptomatic mice (sick) and asymptomatic mice (healthy) from the 28h and 32h groups. (E) miRNAs from the differential analysis performed between mice infected at 8h (early) and mice from the 28h and 32h (late) groups. The common miRNAs between at least two different analyses are indicated by an asterisk (\*).

### Differential expression of host ncRNAs during CDI

ncRNAs can be at the crossroad of regulatory processes governing the interactions of the pathogens with their host during infection [20, 94, 95]. In the present study, many lncRNAs have been identified differentially expressed during CDI, but most of them have not yet been characterized. Compared to uninfected mice, 185 ncRNAs differentially expressed, with more lncRNAs (178 genes, 96 repressed and 82 induced) than microRNAs (7 genes, 4 repressed and 3 induced), have been identified in infected mice. Thirty-eight ncRNAs were differentially expressed in late infected mice, compared to early infected mice (20 induced and 18 repressed), with the vast majority being lncRNAs (36 genes, 20 induced and 16 repressed) and only 2 repressed miRNAs. Among late infected mice, 257 ncRNAs were differentially expressed (155 induced and 102 repressed) in symptomatic mice, compared to asymptomatic mice. As in the global differential analysis encompassing all conditions, almost all of these ncRNAs were lncRNAs (248 genes, 99 repressed and 149 induced). Moreover, in this analysis 9 differentially expressed miRNAs were identified (3 repressed and 6 induced).

Among the ncRNAs, the miRNAs have emerged as important players in host responses to bacterial pathogen infections [94, 96]. We thus extracted the miRNAs differentially expressed in mice during CDI for each comparative analysis (Table 2). Of the 14 miRNAs identified, 7 were repressed and 7 were induced upon CDI. Four miRNAs were found to be differentially expressed in two of the three differential analyses. The two miRNAs, miR-1843b and miR-1949, were repressed in infected mice as compared to uninfected mice but also in late infected mice as compared to early infected mice. miR-1949 has only been described as apoptosis-related miRNA in ovarian granulosa cells induced by cadmium and also as a potential inducer of bladder cancer following spinal cord injury [97, 98]. miR-1938 and miR-3109, were induced in infected mice compared to uninfected mice but also in symptomatic mice compared to asymptomatic mice. These two miRNAs could be then of particular interest for further functional characterization.

Among the miRNAs repressed in mice during CDI, three (miR-145a, miR-682 and miR-99a) have already been involved in anti-inflammatory processes, whereas no role for miR-1843a and miR-3069 has been previously identified. miR-145a negatively regulates the sepsis-induced inflammatory response, through modulation of NF-κB signaling [99], miR-682 has a protective effect on intestinal cells damaged during ischaemic episodes [100] and miR-99a exerts an anti-inflammatory effect when expressed in adipose tissue by inhibiting TNF-α [101].

Among the miRNAs induced in mice during CDI, two (miR-21a and miR-7678) have already been described in inflammatory response. miR-21a, one of the most highly expressed miRNAs in mammalian cells, could play a dynamic role in pro-inflammatory responses [102]. miR-7678 is regulated by TNF-α and involved in controlling the inflammatory response in tissue-engineered cartilage [103]. No role in inflammatory response has been shown for miR-3064 and miR-6236, which have only been described in cardiac or brain diseases [104, 105].

Our results underline the complexity of the regulatory networks of the inflammatory response during CDI and the potential role of miRNAs and lncRNAs in this process. Nevertheless, global miRNA regulation seems to favour the inflammatory process, with reduced expression of anti-inflammatory miRNAs and induction of pro-inflammatory miRNAs. Interestingly, the comparison showing the greatest differences in expression within these regulatory RNAs was between late-infected mice that were sick or asymptomatic. A more detailed analysis of these phenomena could provide a better understanding of the relationship between infection and disease in the host.

## Conclusion

*C. difficile* interacts with host and resident microbial communities inside the gut during infection. We took advantage here of the conventional mouse model of CDI mimicking the infection in humans to follow simultaneously the transcriptome dynamics of the pathogen and the host but also the kinetics of the gut microbiota composition. Such dual *in vivo* transcriptomics approach has never been applied to *C. difficile* before and ncRNAs were not included in previous transcriptomics during CDI, although a recent paper reports a transcriptomic profiling of *C. difficile* attached to epithelial cells using an *in vitro* human gut model over 24h [106]. This study did not look at ncRNA and, importantly, the expression of several key RNA regulators could be only detected under relevant conditions *in vivo*.

From the pathogen side, our data confirmed differential expression *in vivo* as compared to *in vitro* conditions of the toxin, metabolism and sporulation genes also observed with different infection models before and identified for the first time the ncRNA expression dynamics *in vivo*. Our dual RNA-seq analysis revealed new promising candidates among ncRNAs highly induced or repressed *in vivo* that correlated with analysis of available raw RNA-seq datasets from two independent studies. Some of these ncRNAs could be related to the regulation of sporulation process in accordance with accumulating evidence for the importance of RNA-based mechanisms in the control of this key step in *C. difficile* infection cycle including Hfq [17, 107] and Hfq-binding ncRNAs [18, 108, 109].

From the host side, our transcriptomics revealed various inflammation-related pathways as highly induced during infection. A number of known pro-inflammatory miRNAs or previously uncharacterized miRNAs and lncRNAs have been identified as differentially expressed during CDI paving the way for further functional studies of these RNA-based mechanisms modulating host responses. We identified a particular expression pattern for *C. difficile-*infected mice presenting symptoms as compared to infected but asymptomatic mice leading to identification of promising markers associated with extensive inflammatory processes. Unfortunately, the relatively low number of *C. difficile* reads in *in vivo* samples did not allow a detailed comparison of the gene expression profiles between asymptomatic and sick late-infected mice. However, the host changes between these two groups correlated with specific modifications of microbiota profiles revealing interesting candidate species that may be involved in the modulation of the inflammatory process during CDI as potential targets for further microbiota-related modulatory strategies to improve the efficiency of CDI treatments.

Overall, the data generated during this work represent a unique resource for scientific community to explore both the pathogen and the host gene expression during infection and show the power of combined computational approaches applied to complex datasets to extract valuable information on host and pathogen transcriptome, microbiome and ncRNA identification. These data constitute the essential basis to specify the RNA-based mechanisms shaping virulence and adaptation of *C. difficile* to its host and modulating the immune and inflammatory host responses. By identifying specific virulence markers and potential therapeutic targets this work opens new avenues for future development of alternative therapeutic and diagnostic strategies.

## Acknowledgments

This work was supported by grants from the Agence Nationale de la Recherche (“CloSTARn”, ANR-13-JSV3-0005-01, and “CdiffRib”, ANR-22-CE15-0020-01 to O.S.), the Institut Universitaire de France (to O.S.), the University Paris-Saclay, the Institute for Integrative Biology of the Cell, the DIM-1HEALTH regional Ile-de-France program (LSP grant no. 173403, V.K. PhD fellowship). The bioinformatics analyses were performed on the Core Cluster of the Institut Français de Bioinformatique (IFB) (ANR-11-INBS-0013). We are thankful to Pierre Pericard for help in initial dual RNA-seq data analysis.

## Supplementary material

**Figure S1. Clinical signs of sickness indicating *C. difficile* infection for mice during clinical follow-up assay.** Mice were evaluated for stool characteristics, behavior change and weight loss. Each parameter was scored from 0 (formed stool, normal behavior and no change in weight) to 4 (inability to deambulate, mucous stool and weight loss > 15%). Individual scores were combined to a cumulative clinical sickness score (CSS) ranging from 0 to 12.

**Figure S2. Mapping of dual RNA-seq reads against reference genomes of *C. difficile* and mouse and remaining reads of microbiota species in mouse samples.** The total number of read counts mapped on *C. difficile* **(A)** and mouse **(B)** genome, as well as remaining read counts for microbiota species **(C)** are shown for each *in vivo* sample from S1 to S9 for infected mice 8h (in green), 28h (in yellow) and 32h (in red) post-infection as compared to *in vitro* culture samples from IV1 to IV3 (in blue in **(A)**) or uninfected mice samples S10-S12 (in blue in **(B)** and **(C)**).

**Figure S3. Principal component analysis of the differential expression analysis of *C. difficile* genes between groups IV (*in vitro*) and MI (infected mice).** Each *in vivo* sample from S4 to S9 for late-infected mice 28h and 32h post-infection and *in vitro* culture samples (IV) is represented. The PC1 axis allows to separate the samples from the different conditions compared (*in vivo vs in vitro*), meaning that biological variability is the main source of variation in the data analyzed.

**Figure S4. Analysis of microbiota species identified in the mouse gut. (A) Two-dimensional PCA loading plot**. Arrows and text represent the loadings (species) with their contribution on sample separation. **(B) Heatmap of clustering based on species relative abundance across the different samples.** Row scaling was done by applying the unit variance scaling and the clustering of both rows and columns was performed using correlation distance and average linkage. Scale of relative distances is represented on top right ranging from 2 to −2. NOT_INF: mice not infected with *C. difficile* (samples S10, S11, S12), SYMPTOMS: mice infected with *C. difficile* showing visible symptoms (samples S4, S7, S9), NO_SYMPTOMS: mice infected with *C. difficile* showing no visible symptoms (samples S5, S6, S8).

**Figure S5. Relative community composition of the mouse gut microbiota at the species/mOUTs (A) and genus (B) level determined by metatranscriptomics profiling on housekeeping marker genes using mOTUs2.** The experimental conditions are shown under the stacked bars: NOT_INF: mice not infected with *C. difficile* (samples S10-S12). SYMPTOMS: mice infected with *C. difficile* showing visible symptoms (samples S4, S7, S9). NO_SYMPTOMS: mice infected with *C. difficile* showing no visible symptoms (samples S5, S6, S8).

**Figure S6. Functional GSEA analysis of *C. difficile* genes differentially expressed in mice during infection as compared to *in vitro* conditions. (A)** running sum of gsea analysis for Sporulation (left) and Regulations (right) classes; **(B)** proportion of *C. difficile* leading genes given by the gsea analysis with the MA2HTML categories, up (left) and down (right) regulated genes in the mice infected *versus in vitro* growth conditions.

**Figure S7. Real time qRT-PCR of *in vitro*, 28h post-infection (left) and 32h post-infection (right) samples, on *tcdA* (A), and ncRNA SQ995 (B) and SQ1296 (C) genes.** Not infected mice have been used as control. The asterisk indicates a statistically significant difference (* *p*<0.05, ** *p*<0.01, \*\*\**p*<0.001, \*\*\**p*<0.0001). Data are the mean (±SEM) of at least three independent assays.

**Figure S8. Visualization of dual RNA-seq data for *C. difficile* differentially expressed genes with IGV.** Representative examples from different functional groups of genes are presented in **(A)** for protein encoding genes including sporulation proteins, in **(B)** for ncRNAs. The results for *in vivo* samples 28h post-infection (28h pi), 32h post-infection (32h pi) are compared with the data from *in vitro* sample. The genomic regions for previously identified ncRNAs are presented in a green box, newly identified transcribed region are shown in orange box. Genes from MaGe annotation are shown at the bottom of each panel. In the IGV visualization profiles, “+” strand reads are shown in red and “-“ strand reads are shown in blue. IGV visualization is presented with adjusted read threshold for each window to compare the data from different samples (scale is indicated as a read threshold range).

**Figure S9. First two components of a Principal Component Analysis with percentages of variance associated with each axis.** Comparison was made between late infected mice at 28h-32h post-infection (MI) and uninfected control mice (MC) in (**A**); between late infected mice 28h-32h and early infected mice (8h) post-infection (28h-32h) in (**B**) and between symptomatic (sick) and asymptomatic (healthy) mice from the two late infected groups at 28h and 32h post-infection in (**C**).

**Figure S10. Differential Gene Analysis of mouse genes during infection between different groups. (A)** late infected mice at 28h-32h post-infection (MI) were compared to uninfected control mice (MC); **(B)** mice infected at 28h and 32h post-infection (late) were compared to mice infected at 8h post-infection (early); **(C)** symptomatic (sick) mice were compared with asymptomatic (healthy) mice at 28h and 32h post-infection.

**Table S1. Quantification of *C. difficile* vegetative cells in caecal content for each infected mouse.**

**Table S2. Differential Expression Analysis of *C. difficile* genes between late infectious conditions (28h and 32h post-infection) and *in vitro* cultures (excel format in a separate file).**

**Table S3. Enrichment analysis of Ma2HTML classes with *C. difficile* expression profiles in infected mice *versus in vitro* growth conditions.** Term : class name; nes : normalized enrich score; pval : adjusted *p*-value; fdr : false discovery rate; leading_edge/geneset_size : number of leading genes / gene-set size

**Table S4. List of links for downloading of RNA-seq data from previously reported studies.** (*) add the SSR_Id value followed by "_1.fastq.gz" and then "_2.fastq.gz" to get the complete links for downloading of the 2 files of the paired-end sequencing.

**Table S5. Comparison with available *C. difficile in vivo* transcriptomic data (excel format in a separate file).**

**Table S6. Prediction of new transcripts from RNA-seq data with DETR’PROK (excel format in separate file)**

**Table S7. Functional analysis of host genes induced during CDI.** Genes overexpressed in infected mice (MI32H) compared to uninfected control mice (MC8H) (fc, fold change, > 2; p, *p*- value, <0.05; Cluster, cluster number as indicated in Figure 6, the fold changes are those between 32h-infected mice and uninfected mice, but similar results were also obtained between 28h-infected mice and uninfected mice).

**Table S8. Functional analysis of host genes repressed during CDI.** Genes repressed in infected mice (MI32H) compared to uninfected control mice (MC8H) (fc, fold change, < −2; p, *p*-value, <0.05; Cluster, cluster number as indicated in Figure 6)

## References

1. Di Bella S, Sanson G, Monticelli J, Zerbato V, Principe L, Giuffrè M, et al. *Clostridioides difficile* infection: history, epidemiology, risk factors, prevention, clinical manifestations, treatment, and future options. Clin Microbiol Rev 2024; 37: e00135–23.

2. Janoir C. Virulence factors of *Clostridium difficile* and their role during infection. Anaerobe 2016; 37: 13–24.

3. Smits WK, Lyras D, Lacy DB, Wilcox MH, Kuijper EJ. Clostridium difficile infection. Nat Rev Dis Primers 2016; 2: 16020.

4. Batah J, Denève-Larrazet C, Jolivot P-A, Kuehne S, Collignon A, Marvaud J-C, et al. *Clostridium difficile* flagella predominantly activate TLR5-linked NF-κB pathway in epithelial cells. Anaerobe 2016; 38: 116–124.

5. Hör J, Gorski SA, Vogel J. Bacterial RNA Biology on a Genome Scale. Mol Cell 2018; 70: 785–799.

6. Dersch P, Khan MA, Mühlen S, Görke B. Roles of Regulatory RNAs for Antibiotic Resistance in Bacteria and Their Potential Value as Novel Drug Targets. Frontiers in Microbiology 2017; 8.

7. Papenfort K, Vogel J. Regulatory RNA in Bacterial Pathogens. Cell Host & Microbe 2010; 8: 116–127.

8. Soutourina OA, Monot M, Boudry P, Saujet L, Pichon C, Sismeiro O, et al. Genome-Wide Identification of Regulatory RNAs in the Human Pathogen *Clostridium difficile*. PLoS Genet 2013; 9: e1003493.

9. Soutourina O. RNA-based control mechanisms of *Clostridium difficile*. Current Opinion in Microbiology 2017; 36: 62–68.

10. Lamm-Schmidt V, Fuchs M, Sulzer J, Gerovac M, Hör J, Dersch P, et al. Grad-seq identifies KhpB as a global RNA-binding protein in *Clostridioides difficile* that regulates toxin production. microLife 2021; 2: uqab004.

11. Purcell EB, Tamayo R. Cyclic diguanylate signaling in Gram-positive bacteria. FEMS Microbiol Rev 2016; 40: 753–773.

12. Boudry P, Semenova E, Monot M, Datsenko KA, Lopatina A, Sekulovic O, et al. Function of the CRISPR-Cas System of the Human Pathogen *Clostridium difficile*. mBio 2015; 6.

13. Maikova A, Boudry P, Shiriaeva A, Vasileva A, Boutserin A, Medvedeva S, et al. Protospacer-Adjacent Motif Specificity during *Clostridioides difficile* Type I-B CRISPR- Cas Interference and Adaptation. mBio 2021; 12.

14. Maikova A, Severinov K, Soutourina O. New Insights Into Functions and Possible Applications of *Clostridium difficile* CRISPR-Cas System. Front Microbiol 2018; 9: 1740.

15. Soutourina O. Type I Toxin-Antitoxin Systems in Clostridia. Toxins 2019; 11: 253.

16. Peltier J, Hamiot A, Garneau JR, Boudry P, Maikova A, Hajnsdorf E, et al. Type I toxin- antitoxin systems contribute to the maintenance of mobile genetic elements in Clostridioides difficile. Commun Biol 2020; 3: 718.

17. Boudry P, Gracia C, Monot M, Caillet J, Saujet L, Hajnsdorf E, et al. Pleiotropic Role of the RNA Chaperone Protein Hfq in the Human Pathogen *Clostridium difficile*. J Bacteriol 2014; 196: 3234–3248.

18. Boudry P, Piattelli E, Drouineau E, Peltier J, Boutserin A, Lejars M, et al. Identification of RNAs bound by Hfq reveals widespread RNA partners and a sporulation regulator in the human pathogen *Clostridioides difficile*. RNA Biology 2021; 1–22.

19. Fuchs M, Lamm-Schmidt V, Sulzer J, Ponath F, Jenniches L, Kirk JA, et al. An RNA-centric global view of *Clostridioides difficile* reveals broad activity of Hfq in a clinically important gram-positive bacterium. Proc Natl Acad Sci USA 2021; 118: e2103579118.

20. Duval M, Cossart P, Lebreton A. Mammalian microRNAs and long noncoding RNAs in the host-bacterial pathogen crosstalk. Seminars in Cell & Developmental Biology 2017; 65: 11–19.

21. Westermann AJ, Förstner KU, Amman F, Barquist L, Chao Y, Schulte LN, et al. Dual RNA- seq unveils noncoding RNA functions in host–pathogen interactions. Nature 2016; 529: 496–501.

22. Damron FH, Oglesby-Sherrouse AG, Wilks A, Barbier M. Dual-seq transcriptomics reveals the battle for iron during *Pseudomonas aeruginosa* acute murine pneumonia. Sci Rep 2016; 6: 39172.

23. Nuss AM, Beckstette M, Pimenova M, Schmühl C, Opitz W, Pisano F, et al. Tissue dual RNA-seq allows fast discovery of infection-specific functions and riboregulators shaping host–pathogen transcriptomes. Proc Natl Acad Sci USA 2017; 114: E791–E800.

24. Westermann AJ, Barquist L, Vogel J. Resolving host–pathogen interactions by dual RNA- seq. PLoS Pathog 2017; 13: e1006033.

25. Burgess DJ. Host–pathogen duels revealed by dual RNA-seq in vivo. Nature Reviews Genetics 2017; 18: 143–143.

26. Janoir C, Denève C, Bouttier S, Barbut F, Hoys S, Caleechum L, et al. Adaptive Strategies and Pathogenesis of Clostridium difficile from In Vivo Transcriptomics. Infect Immun 2013; 81: 3757–3769.

27. Fletcher JR, Pike CM, Parsons RJ, Rivera AJ, Foley MH, McLaren MR, et al. *Clostridioides difficile* exploits toxin-mediated inflammation to alter the host nutritional landscape and exclude competitors from the gut microbiota. Nat Commun 2021; 12: 462.

28. Fletcher JR, Erwin S, Lanzas C, Theriot CM. Shifts in the Gut Metabolome and Clostridium difficile Transcriptome throughout Colonization and Infection in a Mouse Model. mSphere 2018; 3.

29. Kansau I, Barketi-Klai A, Monot M, Hoys S, Dupuy B, Janoir C, et al. Deciphering Adaptation Strategies of the Epidemic Clostridium difficile 027 Strain during Infection through In Vivo Transcriptional Analysis. PLoS ONE 2016; 11: e0158204.

30. Jenior ML, Leslie JL, Young VB, Schloss PD. *Clostridium difficile* Colonizes Alternative Nutrient Niches during Infection across Distinct Murine Gut Microbiomes. mSystems 2017; 2: e00063–17.

31. Hussain HA, Roberts AP, Mullany P. Generation of an erythromycin-sensitive derivative of *Clostridium difficile* strain 630 (630Δ*erm*) and demonstration that the conjugative transposon Tn916ΔE enters the genome of this strain at multiple sites. Journal of Medical Microbiology 2005; 54: 137–141.

32. Pruss KM, Sonnenburg JL. *C. difficile* exploits a host metabolite produced during toxin-mediated disease. Nature 2021; 593: 261–265.

33. Burns DA, Heeg D, Cartman ST, Minton NP. Reconsidering the Sporulation Characteristics of Hypervirulent *Clostridium difficile* BI/NAP1/027. PLoS ONE 2011; 6: e24894.

34. Chen X, Katchar K, Goldsmith JD, Nanthakumar N, Cheknis A, Gerding DN, et al. A Mouse Model of *Clostridium difficile*–Associated Disease. Gastroenterology 2008; 135: 1984–1992.

35. André G, Even S, Putzer H, Burguière P, Croux C, Danchin A, et al. S-box and T-box riboswitches and antisense RNA control a sulfur metabolic operon of *Clostridium acetobutylicum*. Nucleic Acids Research 2008; 36: 5955–5969.

36. Love MI, Huber W, Anders S. Moderated estimation of fold change and dispersion for RNA-seq data with DESeq2. Genome Biol 2014; 15: 550.

37. Varet H, Brillet-Guéguen L, Coppée J-Y, Dillies M-A. SARTools: A DESeq2- and EdgeR- Based R Pipeline for Comprehensive Differential Analysis of RNA-Seq Data. PLoS ONE 2016; 11: e0157022.

38. R Core Team (2022). R: A language and environment for statistical computing. R Foundation for Statistical Computing, Vienna, Austria. URL https://www.R-project.org/. [2] RStudio Team (2020). RStudio: Integrated Development for R. RStudio, PBC, Boston, MA URL http://www.rstudio.com/.

39. RStudio Team (2020). RStudio: Integrated Development for R. RStudio, PBC, Boston, MA URL http://www.rstudio.com/.

40. Chen Y, Lun ATL, Smyth GK. From reads to genes to pathways: differential expression analysis of RNA-Seq experiments using Rsubread and the edgeR quasi-likelihood pipeline. F1000Res 2016; 5: 1438.

41. Ritchie ME, Phipson B, Wu D, Hu Y, Law CW, Shi W, et al. limma powers differential expression analyses for RNA-sequencing and microarray studies. Nucleic Acids Res 2015; 43: e47.

42. Liberzon A, Birger C, Thorvaldsdóttir H, Ghandi M, Mesirov JP, Tamayo P. The Molecular Signatures Database (MSigDB) hallmark gene set collection. Cell Syst 2015; 1: 417–425.

43. Monot M, Orgeur M, Camiade E, Brehier C, Dupuy B. COV2HTML: A Visualization and Analysis Tool of Bacterial Next Generation Sequencing (NGS) Data for Postgenomics Life Scientists. OMICS: A Journal of Integrative Biology 2014; 18: 184–195.

44. Lachmann A, Xie Z, Ma’ayan A. blitzGSEA: efficient computation of gene set enrichment analysis through gamma distribution approximation. Bioinformatics 2022; 38: 2356–2357.

45. Milanese A, Mende DR, Paoli L, Salazar G, Ruscheweyh H-J, Cuenca M, et al. Microbial abundance, activity and population genomic profiling with mOTUs2. Nat Commun 2019; 10: 1014.

46. Dobin A, Davis CA, Schlesinger F, Drenkow J, Zaleski C, Jha S, et al. STAR: ultrafast universal RNA-seq aligner. Bioinformatics 2013; 29: 15–21.

47. Langmead B, Salzberg SL. Fast gapped-read alignment with Bowtie 2. Nat Methods 2012; 9: 357–359.

48. Segata N, Izard J, Waldron L, Gevers D, Miropolsky L, Garrett WS, et al. Metagenomic biomarker discovery and explanation. Genome Biol 2011; 12: R60.

49. Köster J, Rahmann S. Snakemake—a scalable bioinformatics workflow engine. Bioinformatics 2012; 28: 2520–2522.

50. Larsson J, Gustafsson P. A Case Study in Fitting Area-Proportional Euler Diagrams with Ellipses using eulerr. 2018.

51. Gu Z, Eils R, Schlesner M. Complex heatmaps reveal patterns and correlations in multidimensional genomic data. Bioinformatics 2016; 32: 2847–2849.

52. Toffano-Nioche C, Luo Y, Kuchly C, Wallon C, Steinbach D, Zytnicki M, et al. Detection of non-coding RNA in bacteria and archaea using the DETR’PROK Galaxy pipeline. Methods 2013; 63: 60–65.

53. Zytnicki M, Quesneville H. S-MART, A Software Toolbox to Aid RNA-seq Data Analysis. PLoS ONE 2011; 6: e25988.

54. D’Auria KM, Kolling GL, Donato GM, Warren CA, Gray MC, Hewlett EL, et al. In Vivo Physiological and Transcriptional Profiling Reveals Host Responses to Clostridium difficile Toxin A and Toxin B. Infect Immun 2013; 81: 3814–3824.

55. Passmore IJ, Letertre MPM, Preston MD, Bianconi I, Harrison MA, Nasher F, et al. Para- cresol production by *Clostridium difficile* affects microbial diversity and membrane integrity of Gram-negative bacteria. PLoS Pathog 2018; 14: e1007191.

56. Darkoh C, DuPont HL, Norris SJ, Kaplan HB. Toxin Synthesis by *Clostridium difficile* Is Regulated through Quorum Signaling. mBio 2015; 6.

57. Theriot CM, Young VB. Interactions Between the Gastrointestinal Microbiome and *Clostridium difficile*. Annu Rev Microbiol 2015; 69: 445–461.

58. Pike CM, Theriot CM. Mechanisms of Colonization Resistance Against *Clostridioides difficile*. J Infect Dis 2021; 223: S194–S200.

59. Martinez E, Taminiau B, Rodriguez C, Daube G. Gut Microbiota Composition Associated with *Clostridioides difficile* Colonization and Infection. Pathogens 2022; 11: 781.

60. Schubert AM, Rogers MAM, Ring C, Mogle J, Petrosino JP, Young VB, et al. Microbiome data distinguish patients with *Clostridium difficile* infection and non-*C. difficile*- associated diarrhea from healthy controls. mBio 2014; 5: e01021–01014.

61. Spinler JK, Auchtung J, Brown A, Boonma P, Oezguen N, Ross CL, et al. Next-Generation Probiotics Targeting *Clostridium difficile* through Precursor-Directed Antimicrobial Biosynthesis. Infect Immun 2017; 85: e00303–17.

62. Quigley L, Coakley M, Alemayehu D, Rea MC, Casey PG, O’Sullivan Ó, et al. Lactobacillus gasseri APC 678 Reduces Shedding of the Pathogen *Clostridium difficile* in a Murine Model. Front Microbiol 2019; 10: 273.

63. Charlet R, Le Danvic C, Sendid B, Nagnan-Le Meillour P, Jawhara S. Oleic Acid and Palmitic Acid from *Bacteroides thetaiotaomicron* and *Lactobacillus johnsonii* Exhibit Anti-Inflammatory and Antifungal Properties. Microorganisms 2022; 10: 1803.

64. Schubert AM, Sinani H, Schloss PD. Antibiotic-Induced Alterations of the Murine Gut Microbiota and Subsequent Effects on Colonization Resistance against *Clostridium difficile*. mBio 2015; 6: e00974.

65. Hamilton MJ, Weingarden AR, Unno T, Khoruts A, Sadowsky MJ. High-throughput DNA sequence analysis reveals stable engraftment of gut microbiota following transplantation of previously frozen fecal bacteria. Gut Microbes 2013; 4: 125–135.

66. Goldberg E, Amir I, Zafran M, Gophna U, Samra Z, Pitlik S, et al. The correlation between *Clostridium-difficile* infection and human gut concentrations of Bacteroidetes phylum and clostridial species. Eur J Clin Microbiol Infect Dis 2014; 33: 377–383.

67. Deng H, Yang S, Zhang Y, Qian K, Zhang Z, Liu Y, et al. Bacteroides fragilis Prevents *Clostridium difficile* Infection in a Mouse Model by Restoring Gut Barrier and Microbiome Regulation. Front Microbiol 2018; 9: 2976.

68. Ghimire S, Roy C, Wongkuna S, Antony L, Maji A, Keena MC, et al. Identification of *Clostridioides difficile*-Inhibiting Gut Commensals Using Culturomics, Phenotyping, and Combinatorial Community Assembly. mSystems 2020; 5: e00620–19.

69. Satokari R, Fuentes S, Mattila E, Jalanka J, de Vos WM, Arkkila P. Fecal transplantation treatment of antibiotic-induced, noninfectious colitis and long-term microbiota follow- up. Case Rep Med 2014; 2014: 913867.

70. Sekulovic O, Ospina Bedoya M, Fivian-Hughes AS, Fairweather NF, Fortier L. The *Clostridium difficile* cell wall protein CWPV confers phase-variable phage resistance. Molecular Microbiology 2015; 98: 329–342.

71. Reynolds CB, Emerson JE, de la Riva L, Fagan RP, Fairweather NF. The *Clostridium difficile* cell wall protein CwpV is antigenically variable between strains, but exhibits conserved aggregation-promoting function. PLoS Pathog 2011; 7: e1002024–e1002024.

72. Jenior ML, Leslie JL, Powers DA, Garrett EM, Walker KA, Dickenson ME, et al. Novel Drivers of Virulence in *Clostridioides difficile* Identified via Context-Specific Metabolic Network Analysis. mSystems 2021; 6: e0091921.

73. Scaria J, Janvilisri T, Fubini S, Gleed RD, McDonough SP, Chang Y-F. *Clostridium difficile* Transcriptome Analysis Using Pig Ligated Loop Model Reveals Modulation of Pathways Not Modulated In Vitro. The Journal of Infectious Diseases 2011; 203: 1613–1620.

74. Ternan NG, Jain S, Srivastava M, McMullan G. Comparative transcriptional analysis of clinically relevant heat stress response in *Clostridium difficile* strain 630. PLoS One 2012; 7: e42410.

75. Purcell EB, McKee RW, McBride SM, Waters CM, Tamayo R. Cyclic Diguanylate Inversely Regulates Motility and Aggregation in *Clostridium difficile*. J Bacteriol 2012; 194: 3307– 3316.

76. Bordeleau E, Burrus V. Cyclic-di-GMP signaling in the Gram-positive pathogen *Clostridium difficile*. Curr Genet 2015; 61: 497–502.

77. Battaglioli EJ, Hale VL, Chen J, Jeraldo P, Ruiz-Mojica C, Schmidt BA, et al. *Clostridioides difficile* uses amino acids associated with gut microbial dysbiosis in a subset of patients with diarrhea. Sci Transl Med 2018; 10: eaam7019.

78. Reed AD, Fletcher JR, Huang YY, Thanissery R, Rivera AJ, Parsons RJ, et al. The Stickland Reaction Precursor trans -4-Hydroxy- L -Proline Differentially Impacts the Metabolism of Clostridioides difficile and Commensal Clostridia. mSphere 2022; 7: e00926-21.

79. Ho TD, Ellermeier CD. Ferric Uptake Regulator Fur Control of Putative Iron Acquisition Systems in *Clostridium difficile*. J Bacteriol 2015; 197: 2930–2940.

80. Hastie JL, Hanna PC, Carlson PE. Transcriptional response of *Clostridium difficile* to low iron conditions. Pathog Dis 2018; 76: fty009.

81. Berges M, Michel A-M, Lassek C, Nuss AM, Beckstette M, Dersch P, et al. Iron Regulation in *Clostridioides difficile*. Front Microbiol 2018; 9: 3183.

82. Saujet L, Pereira FC, Serrano M, Soutourina O, Monot M, Shelyakin PV, et al. Genome- Wide Analysis of Cell Type-Specific Gene Transcription during Spore Formation in *Clostridium difficile*. PLoS Genet 2013; 9: e1003756.

83. Fimlaid KA, Bond JP, Schutz KC, Putnam EE, Leung JM, Lawley TD, et al. Global Analysis of the Sporulation Pathway of *Clostridium difficile*. PLoS Genet 2013; 9: e1003660.

84. Soutourina O, Dubois T, Monot M, Shelyakin PV, Saujet L, Boudry P, et al. Genome-Wide Transcription Start Site Mapping and Promoter Assignments to a Sigma Factor in the Human Enteropathogen *Clostridioides difficile*. Front Microbiol 2020; 11: 1939.

85. Kreis V, Soutourina O. *Clostridioides difficile* – phage relationship the RNA way. Current Opinion in Microbiology 2022; 66: 1–10.

86. R Andrade P, Mehta M, Lu J, M B Teles R, Montoya D, O Scumpia P, et al. The cell fate regulator NUPR1 is induced by *Mycobacterium leprae* via type I interferon in human leprosy. PLoS Negl Trop Dis 2019; 13: e0007589.

87. Rajamani D, Singh PK, Rottmann BG, Singh N, Bhasin MK, Kumar A. Temporal retinal transcriptome and systems biology analysis identifies key pathways and hub genes in *Staphylococcus aureus* endophthalmitis. Sci Rep 2016; 6: 21502.

88. Rosengren AT, Nyman TA, Lahesmaa R. Proteome profiling of interleukin-12 treated human T helper cells. Proteomics 2005; 5: 3137–3141.

89. Xi R, Zheng X, Tizzano M. Role of Taste Receptors in Innate Immunity and Oral Health. J Dent Res 2022; 101: 759–768.

90. Sandholt AKS, Wattrang E, Lilja T, Ahola H, Lundén A, Troell K, et al. Dual RNA-seq transcriptome analysis of caecal tissue during primary *Eimeria tenella* infection in chickens. BMC Genomics 2021; 22: 660.

91. Wesener DA, Wangkanont K, McBride R, Song X, Kraft MB, Hodges HL, et al. Recognition of microbial glycans by human intelectin-1. Nat Struct Mol Biol 2015; 22: 603–610.

92. Sawant M, Benamrouz-Vanneste S, Mouray A, Bouquet P, Gantois N, Creusy C, et al. Persistent Cryptosporidium parvum Infection Leads to the Development of the Tumor Microenvironment in an Experimental Mouse Model: Results of a Microarray Approach. Microorganisms 2021; 9: 2569.

93. Gillespie M, Jassal B, Stephan R, Milacic M, Rothfels K, Senff-Ribeiro A, et al. The reactome pathway knowledgebase 2022. Nucleic Acids Research 2022; 50: D687–D692.

94. Riahi Rad Z, Riahi Rad Z, Goudarzi H, Goudarzi M, Mahmoudi M, Yasbolaghi Sharahi J, et al. MicroRNAs in the interaction between host-bacterial pathogens: A new perspective. J Cell Physiol 2021; 236: 6249–6270.

95. Schmerer N, Schulte LN. Long noncoding RNAs in bacterial infection. Wiley Interdiscip Rev RNA 2021; 12: e1664.

96. Aguilar C, Mano M, Eulalio A. MicroRNAs at the Host-Bacteria Interface: Host Defense or Bacterial Offense. Trends Microbiol 2019; 27: 206–218.

97. Sun Y, Lv Y, Li Y, Li J, Liu J, Luo L, et al. Maternal genetic effect on apoptosis of ovarian granulosa cells induced by cadmium. Food Chem Toxicol 2022; 165: 113079.

98. Wang T, Liu Y, Yuan W, Zhang L, Zhang Y, Wang Z, et al. Identification of microRNAome in rat bladder reveals miR-1949 as a potential inducer of bladder cancer following spinal cord injury. Mol Med Rep 2015; 12: 2849–2857.

99. Wu Y, Li P, Goodwin AJ, Cook JA, Halushka PV, Zingarelli B, et al. miR-145a Regulation of Pericyte Dysfunction in a Murine Model of Sepsis. J Infect Dis 2020; 222: 1037–1045.

100. Liu Z, Jiang J, Yang Q, Xiong Y, Zou D, Yang C, et al. MicroRNA-682-mediated downregulation of PTEN in intestinal epithelial cells ameliorates intestinal ischemia- reperfusion injury. Cell Death Dis 2016; 7: e2210.

101. Jaiswal A, Reddy SS, Maurya M, Maurya P, Barthwal MK. MicroRNA-99a mimics inhibit M1 macrophage phenotype and adipose tissue inflammation by targeting TNFα. Cell Mol Immunol 2019; 16: 495–507.

102. Sheedy FJ. Turning 21: Induction of miR-21 as a Key Switch in the Inflammatory Response. Front Immunol 2015; 6: 19.

103. Ross AK, Coutinho de Almeida R, Ramos YFM, Li J, Meulenbelt I, Guilak F. The miRNA- mRNA interactome of murine induced pluripotent stem cell-derived chondrocytes in response to inflammatory cytokines. FASEB J 2020; 34: 11546–11561.

104. Yang W, Tu H, Tang K, Huang H, Ou S, Wu J. MiR-3064 in Epicardial Adipose-Derived Exosomes Targets Neuronatin to Regulate Adipogenic Differentiation of Epicardial Adipose Stem Cells. Front Cardiovasc Med 2021; 8: 709079.

105. Ko J, Hemphill M, Yang Z, Beard K, Sewell E, Shallcross J, et al. Multi-Dimensional Mapping of Brain-Derived Extracellular Vesicle MicroRNA Biomarker for Traumatic Brain Injury Diagnostics. J Neurotrauma 2020; 37: 2424–2434.

106. Frost LR, Stark R, Anonye BO, MacCreath TO, Ferreira LRP, Unnikrishnan M. Dual RNA- seq identifies genes and pathways modulated during *Clostridioides difficile* colonization. mSystems 2023; 8: e00555–23.

107. Maikova A, Kreis V, Boutserin A, Severinov K, Soutourina O. Using an Endogenous CRISPR-Cas System for Genome Editing in the Human Pathogen *Clostridium difficile*. Appl Environ Microbiol 2019; 85.

108. Fuchs M, Lamm-Schmidt V, Lenče T, Sulzer J, Bublitz A, Wackenreuter J, et al. A network of small RNAs regulates sporulation initiation in *Clostridioides difficile*. EMBO J 2023; 42: e112858.

109. Edwards AN, McBride SM. The RgaS-RgaR two-component system promotes *Clostridioides difficile* sporulation through a small RNA and the Agr1 system. PLoS Genet 2023; 19: e1010841.

